# The SIR2/3/4 complex promotes efficient break-induced replication by stabilizing recombination intermediates

**DOI:** 10.1101/2025.05.22.655457

**Authors:** Beata L Mackenroth, Eric Alani

## Abstract

Homologous recombination (HR) is one of the most effective pathways a cell can employ to repair DNA double strand breaks (DSBs). While most of the HR machinery has been characterized, less is known about how chromatin is reorganized and participates in repair. Here, we use a break-induced replication (BIR) assay, which measures conservative copying of a donor chromosome arm onto a single-ended recipient sequence, to explore a direct role for the SIR2/3/4 (SIR) silent chromatin complex in HR. Specifically, we measure an early step of BIR, D-loop extension, and simultaneously monitor localization of the SIR complex at the initiating DSB and an ectopic donor locus. Interestingly, we observe an ∼four-fold increase in Sir2 and Sir3 protein levels upon DSB induction and detect an enrichment of all three SIR complex subunits at the DSB and donor loci during periods of active D-loop extension. ChIP-seq experiments reveal enrichment of SIR complex subunits that extend up to 15 kb from the DSB site. This enrichment is substantially reduced in *rad51Δ* and *rad52Δ* mutants that show defects in BIR. Lastly, we show that *sir* mutations confer slowed D-loop extension kinetics between divergent substrates and increase the rate of mutation during BIR. These data are consistent with a model in which the SIR complex acts to stabilize the extending D-loop during BIR, leading to more efficient, and thus less mutagenic repair.

**Author Summary:** Double-strand breaks (DSBs) are among the most dangerous types of DNA lesions that occur in a cell; if unrepaired, they can cause cell death and deleterious chromosomal rearrangements. Homologous recombination (HR) is one of the most effective pathways to repair DSBs. While the machinery involved in HR has been characterized, less is known about how HR acts within a chromatin environment which contains DNA packaged into a compact state. In baker’s yeast transcriptionally inactive “silent” chromatin is present at a limited number of domains in the cell. Bound to and regulating the transcriptional inactivity of these domains is the SIR complex. Curiously, various components of the SIR complex have been shown to localize to HR repaired DSBs. However, little is known about how they participate in HR. In this study we use a break-induced replication (BIR) assay, which measures conservative copying of a donor chromosome onto a single-ended recipient sequence, to characterize roles for the SIR complex in HR. We detect an enrichment of all three SIR complex subunits at the recipient and donor loci during periods of active repair. ChIP-seq, a method that detects protein-DNA interactions genome-wide, reveals enrichment of SIR complex subunits up to 15 kb from each side of the DSB site. This enrichment is significantly reduced in mutants that show defects in BIR. We assess the functional consequences of the loss of the SIR complex, showing that *sir* mutations slow early steps in BIR and increase the rate of mutation. Our data are consistent with a model in which the SIR complex acts to stabilize early intermediates in BIR, leading to more efficient, and thus less mutagenic repair.

## Introduction

Preserving DNA integrity is critical for cell survival and organismal growth and development. Remarkably, human cells maintain their genome integrity despite incurring an estimated 50 DSBs per cell cycle [1]. One process used to repair DSBs is non-homologous end joining (NHEJ), in which broken DNA ends bound by the Mre11-Rad50-Xrs2 (MRX) and Yku70/80 complexes are ultimately ligated by DNA ligase IV (baker’s yeast nomenclature used here; reviewed in [2]). This repair is not strictly conservative and can lead to insertion/deletion mutations, or gross chromosome rearrangements. A second DSB repair pathway is homologous recombination (HR; reviewed in [2–6]) in which DSB ends are first resected to form 3’ single-stranded ends that are then coated by the Rad51 recombinase [7,8].

The resulting nucleoprotein filament invades the homologous donor locus to form a D-loop structure [9–12] (Fig 1A). This D-loop is extended by DNA Polymerase δ using the donor as a template, and DSB repair can then proceed through several sub-pathways, often resulting in conservative repair [13].

**Fig 1.**
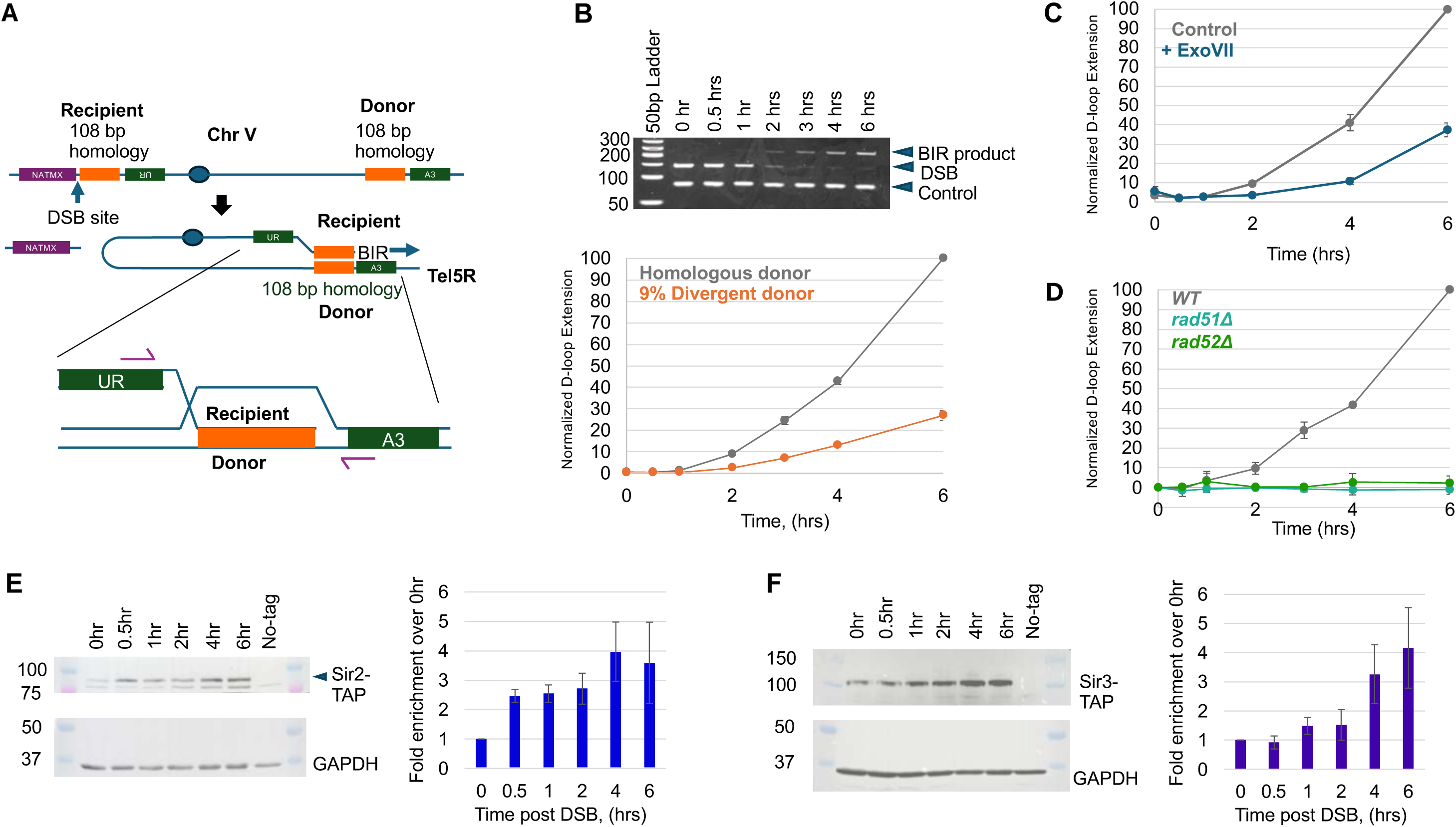
System used to measure repair through BIR. (A) Schematic of BIR system used in this study. A single DSB on Chromosome V is induced by expressing the HO endonuclease through the *GAL1/10* promoter -30 kb from Tel5L ("Recipient" locus) [58]. One side of the DSB has a 108 bp stretch of homology a "Donor" locus, -20 kb from Tel5R. Repair can occur via invasion to form a D-loop structure, and BIR to the end of the chromosome. This results in the formation of a functional *URA3* gene, and the loss of the left arm of Chromosome V, which is marked with *NATMX*. Extension of the D-loop by ∼40 bp can be measured by PCR, using primers specific to the chimeric molecule formed during repair. This is visualized in (B), top panel, alongside primers to measure DSB formation (amplicon is lost upon DSB induction), and control primers targeting *ACT1*. This D-loop extension product is measured by qPCR (B), lower panel, normalized to PCRs to a single copy control locus *ZWF1* on Chromosome VI (Materials and Methods). (C). Sensitivity of D-loop extension qPCR product to degradation by ExoVII nuclease. Control amplicons were treated/prepared in the same manner as ExoVII treated ones, except for the absence of ExoVII nuclease. Error bars represent SEM. (D). D-loop extension product, measured by qPCR, comparing *wild-type, rad51Δ* and *rad52Δ* mutants. Error bars represent SEM. (E, F) Representative Western blots of Sir2-TAP (E), Sir3-TAP (F), 0-6 hrs post DSB induction with galactose. Three independent experiments of independent transformants were quantified and normalized by GAPDH loading controls. Error bars represent SEM.

In some instances, single-ended DSBs may arise through the stalling and subsequent dissolution of replication forks or the inhibition of topoisomerases (reviewed in [14–16]). Lacking a second end, these DSBs are more difficult to repair by the mechanisms described above, and an HR mechanism termed break induced replication (BIR) is often employed [17–20]. During BIR a D-loop is formed (Fig 1A), but in the absence of a second end, DNA Polymerase δ continues to extend from the D-loop.

Synthesis of the complementary DNA strand is achieved by a second DNA polymerase δ which in conjunction with α primase, uses the newly extended strand as a template [21–23]. This process is discontinuous and may lag up to 30 kb from first strand synthesis [24]. Stepwise synthesis is thought to continue to the end of the chromosome, resulting in the fully conservative copying of the donor chromosome arm onto the recipient.

Several steps in BIR have been linked to mutagenesis [25–27]. First, the D-loop is unstable, and the 3’ single-stranded end may undergo several rounds of invasion, possibly using different donor templates each time. Second, the long-lived single-stranded DNA intermediates present during BIR are highly sensitive to damage [28]. Finally, invasion of a non-allelic donor template can lead to loss of heterozygosity across the newly synthesized chromosome arm or gross chromosomal rearrangements.

Notably, in human cancer cells, BIR-like mechanisms are thought to underly patterns of mutagenesis such as kataegis, involving hypermutation at localized regions, and chromothripsis, as well as alternative lengthening of telomeres [27–30]. Thus, understanding BIR, for which baker’s yeast provides an excellent model, can give us insight into HR mechanisms in higher eukaryotes.

To this already complex mechanism is the need to understand how this process occurs in a chromatin environment. Initial efforts to understand these interactions led to the “access, repair, restore” model. This model describes recruitment of deacetylases and chromatin remodelers to damaged DNA to allow access to factors which enact repair, followed by recruitment of chromatin chaperones and deacetylases to restore chromatin to its pre-DSB state. Beyond this general sketch, the role of chromatin in DSB repair appears to differ across repair processes (reviewed in [2,31,32]). In some contexts, chromatin factors play pro-recombination roles such as in fission yeast, where deposition of histones by CAF-1 stabilizes recombination intermediates [33].

In baker’s yeast most of the genome is present in a transcriptionally active “open” chromatin state. The yeast nucleus is spatially organized with the centromeres bound to a peripheral location; similarly, the small amount of transcriptionally inactive “silent” chromatin at the silent mating type loci *HMR* and *HML* and the yeast telomeres form 5-6 bundles at the nuclear periphery [34–36]. Bound to, and regulating the transcriptional inactivity of these domains is the Sir2/Sir3/Sir4 (SIR) complex. The SIR complex is comprised of three subunits: Sir2, a histone deacetylase, Sir3, which contains DNA- and nucleosome-binding domains, and Sir4, which serves as a scaffold, and has several interaction domains. This complex forms a nucleosome-bound 30 nm filament [37,38]. DNA sequences called silencers are bound by several factors such as ORC, Rap1, Abf1, Sir1, and yKu70/80, which serve as nucleation points for SIR filaments [39–43]. SIR complex binding and filament growth are tightly regulated, are thought to occur in a stepwise fashion, and are dependent upon the deacetylase activity of Sir2 [38,44]. Once bound to silent mating type loci or yeast telomeres, the SIR filament participates in tethering these loci to the nuclear periphery through the interaction of Sir4 and Esc1 [43,45,46].

Over the last few decades, the SIR complex has been shown to participate in other nuclear processes, notably DSB repair. First, SIR complex mutants affect the efficiency of non-homologous end joining in yeast [47,48]. One explanation for this phenomenon is that *sir* mutants display a “pseudo diploid” phenotype due to mis-expression of the *MATa* and *MATalpha* genes at *HMR* and *HML;* such a phenotype is associated with a decrease in repair by NHEJ [49]. However, direct roles for the SIR complex in DSB repair have been suggested by the physical interaction of Sir4 with the yKu70/80 complex [47,48,50]; yKu70/80 promotes NHEJ, but has also been shown to stimulate MRX dependent resection of DSBs for repair via HR [51]. Recently, Sir3 has been shown to bind and inhibit Sae2, a pro-resection factor, to promote NHEJ rather than HR [52].

While several studies support roles for subunits of the SIR complex in promoting efficient NHEJ, other work suggests that the SIR complex may act in HR. For instance, Sir2 has been shown to localize to DSBs 2-3 hours post DSB induction during HR mediated mating type switching [53], and Sir3 and Sir4 were found to localize to an unrepairable DSB 4-6 hours post DSB induction [54] . Yet other work showed that in S-phase, where HR repair of DSBs predominates, Sir3 was released from baker’s yeast telomeres and localized to DSBs. Such Sir3 localization was not observed in G1, where baker’s yeast prefers NHEJ [55,56].

A few groups have sought to investigate the role of the SIR complex in HR using genetic assays [53,57]. However, such experiments utilized mating type switching on Chromosome III as a model, where normal binding of the SIR complex at the donor templates *HMR* and *HML* complicates the interpretation of results. In this study, we utilized an inducible single-DSB system that involves repair by BIR [58,59]. Critically, the DSB and donor sequences are not normally bound by the SIR complex. To characterize direct effects of the SIR complex, we examine the recruitment of SIR2/3/4 to the donor and recipient and assess the functional consequences of the loss of the SIR complex in an early step of BIR, D-loop extension. We observe SIR complex binding at recipient and donor loci and the establishment of a large SIR complex bound domain upstream of the DSB locus. This SIR binding tract is dependent upon repair by HR and is altered in the absence of the DNA mismatch repair protein Msh2. Further, loss of SIR factors slows D-loop extension and increases rates of mutagenesis during BIR.

## Results

To directly investigate the role of the SIR complex in HR, we looked for a system where a single DSB could be induced, a D-loop could be formed in a single orientation at a single locus, and repair could proceed relatively efficiently. We take advantage of strains developed by the Haber lab [58], which utilize an HO endonuclease induced DSB on one end of Chromosome V, one side of which possesses homology to an ectopic donor site located on the other end of the chromosome (S1 Table; Fig 1A). We induce a DSB by shifting cells to media containing galactose, thus expressing the HO endonuclease under control of the *GAL1/10* promoter. Repair of this side of the DSB involves invasion at the 108 bp homologous donor sequence followed by conservative DNA synthesis (∼25 kb) by BIR to the end of the chromosome. Such repair results in the formation of a functional *URA3* gene and thus a Ura^+^ phenotype. Critically, in this background the HO endonuclease cut at the *MAT* locus has been mutated such that it is uncleavable, and the silent mating type loci *HMR* and *HML*, which would be de-silenced by mutation of the SIR complex, have been removed. Thus, the absence of the SIR complex in this background does not cause any change in *MATa/alpha* gene expression, a phenomenon which would alter expression of certain recombination factors such as Rdh54 [11,60].

We observe BIR for *wild-type* strains with homologous donors, as measured by Ura^+^ survival, to be 11 +/- 0.5 % (SEM; S1_Data, Underlying Data), comparable to previously published observations [58,59]. In this system, the D-loop must form in a single orientation and extend unidirectionally. This feature, coupled with the short length of donor homology in the strain, allows us to assess by qPCR an early step in BIR, the initiation of new DNA synthesis. (Fig 1B). The efficiency of this reaction is dependent donor homology; when provided a 9% divergent donor sequence, this synthesis, which we refer to as D-loop extension (see below), decreases roughly 3.5-fold (Fig 1B). Importantly, the D-loop extension signal is dependent upon HR; in the absence of key HR proteins Rad51 or Rad52, the signal is lost (Fig 1D).

There has been some question as to whether such qPCR products capture the single-stranded D-loop extension product, or only the double-stranded product formed after second strand synthesis [12,24]. Consistent with qPCR detecting D-loop extension, we observed that the bulk of the substrate amplified by the D-loop extension primers is sensitive to treatment with the single-stranded DNA nuclease ExoVII (Fig 1C). We also note that a single-stranded substrate produces slightly less qPCR signal than a double-stranded one. Thus, our observations are probably an underestimate of the proportion of single-stranded substrate at a given timepoint.

To characterize protein dynamics of the SIR complex during the process of BIR, we tagged Sir2, Sir3, and Sir4 with C-terminal TAP tags from the yeast TAP collection [61,62]. These tags have been used previously for ChIP [63], and found to be functional for silencing [64–66]. In our hands, Sir2, Sir3, and Sir4-TAP strains are haploproficient (S1 Fig), and both Sir2 and Sir3 migrate in crude extracts at their expected sizes (Figs 1E and 1F; S2 Fig). Unfortunately, Sir4 proved difficult to detect via Western blot, likely due to its large size and extensive SUMOylation [67,68]. Upon DSB induction, we observe that both Sir2 and Sir3 protein levels increase 3- to 4-fold within 6 hours (Figs 1E and 1F; S2 Fig). This observation was surprising; SIR protein expression levels are tightly regulated; in fact, previous work showed that overexpression of the SIR subunits can lead to telomere hyperclustering, aberrant extension of SIR filaments into nearby euchromatic regions, and cell death [69,70]. These observations suggest to us that the SIR complex performs some function during BIR.

To investigate a direct role of the SIR complex in BIR, we performed ChIP-qPCR for Sir2, Sir3, and Sir4 around the site of D-loop formation. The ectopic repair of the DSB via BIR allows us to unequivocally ascribe enrichment to donor and recipient sequences (Fig 2A). In our experiments, we take samples relatively early during BIR (up to 6 hours post DSB), because the DNA Damage Response (DDR) checkpoint should prevent these cells from undergoing DNA replication during repair. This is supported by the observations of Liu et al, who do not observe significant differences in BIR kinetics between arrested and non-arrested cells [24]. Intriguingly, we observe significant enrichment of SIR complex subunits, upstream of the extending D-loop, on both recipient and donor strands at 4 and 6 hour timepoints (Figs 2B, 2C, 2E, and 2F; S3A and S3B Figs). This enrichment occurs well into the process of BIR and coincides with the most active periods of D-loop extension (Figs 2D and 2G; S3C Fig). We note variability between replicates, which mirrors the variability observed in the D-loop extension kinetics before wild-type normalization (S4 Fig). We surmise from these data that SIR complex binding is somewhat transient, and that the SIR complex is unlikely to remain bound after repair is complete. We were intrigued that enrichment occurs across a large distance, from 1 to 10 kb.

**Fig 2.**
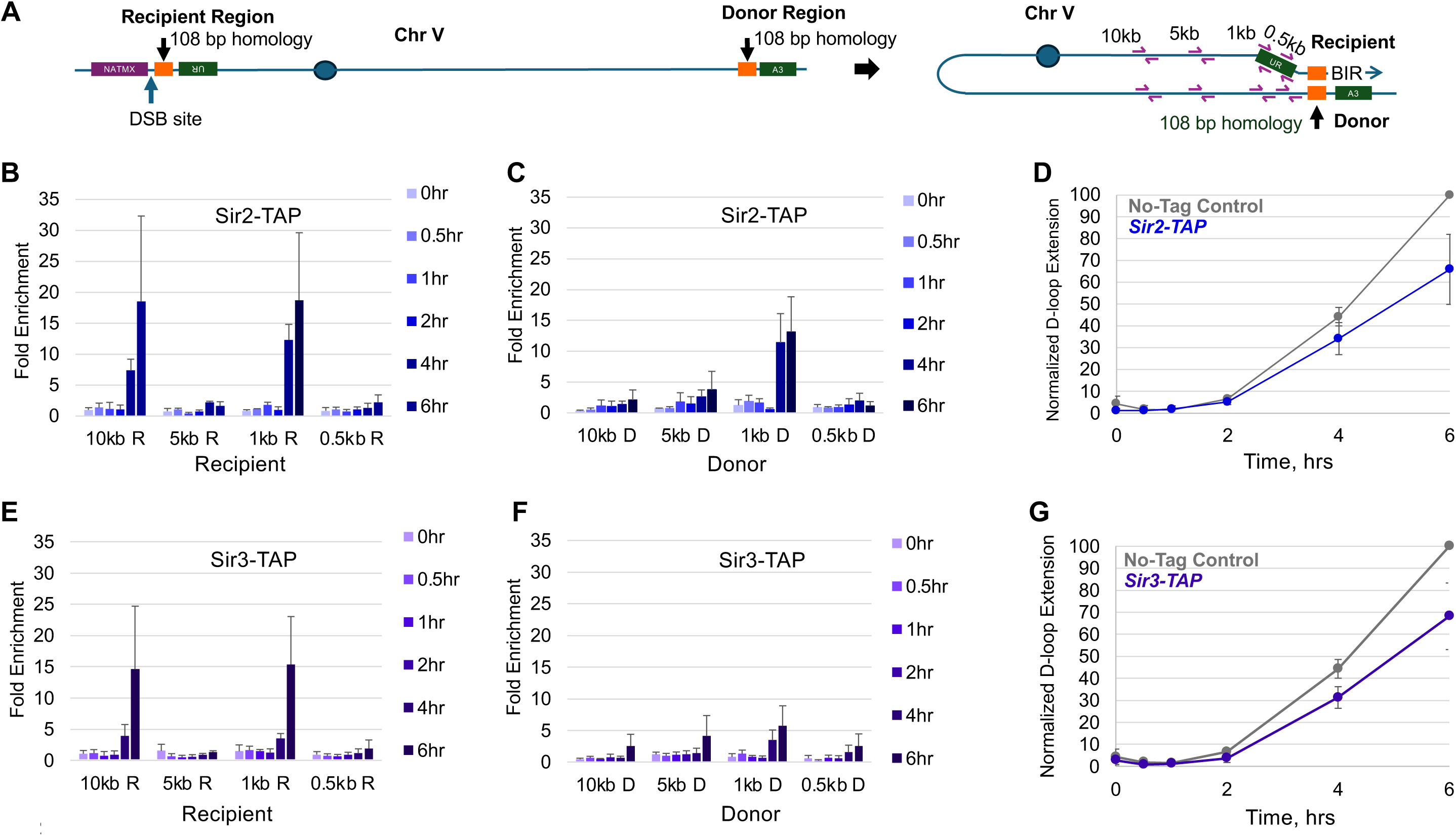
Enrichment of Sir2 and Sir3 at donor and recipient loci during early BIR. (A) Schematic representing qPCR primer location in the ChlP-qPCR analysis. (B, C) Average fold enrichment over No-Tag control strains for Sir2-TAP at Recipient (B) and Donor (C) loci (at 0.5, 1, 5, and 10 kb upstream of each locus). (D) D-loop extension data corresponding to the time courses in (B) and (C). (E, F) Fold enrichment over No-Tag control strains for Sir3-TAP at Recipient (E) and Donor (F) loci. (G) D-loop extension data corresponding to the time courses in (E) and (F). For the ChIP and D-loop extension experiments, averages are the result of three independent transformants, assayed on separate days. Error bars represent SEM.

When comparing both donor and recipient strands, we note that the magnitude of enrichment is generally greater at the recipient than at the donor (Fig 2). This is consistent with the observation that nearly all cells receive a DSB, and in exponentially growing yeast, where most cells are in S-phase, repair via HR is greatly favored [2]. However, while most of the cells can be expected to attempt repair via HR, a far lower percentage would be expected to successfully find and utilize the small ectopic donor locus, as evidenced by BIR having an ∼10 to 15% successful rate [58]. We also note variability in enrichment across the upstream loci (Fig 2; S3 Fig). This is most likely a result of the methods employed during our ChIP workflow. Solubilizing large, heterochromatic filaments can be difficult, and we quickly discovered that the sonication necessary to solubilize the SIR complex subunits resulted in significant protein degradation, even when samples were continuously chilled. This is consistent with observations that sonication specifically degrades large, rigid structures, while having less effect on smaller proteins [71]. Thus, we turned to treatment with double-strand DNA nucleases to solubilize chromatin without protein degradation (Materials and Methods). This was necessary to preserve protein integrity but had the effect of “footprinting” the DNA input to the size of a nucleosome fragment, and may not have solubilized tightly bound, well established SIR filaments. Thus, successful detection of SIR complex enrichment via ChIP-qPCR requires primers to lie fully within a ∼150 bp nucleosome fragment, and well-established filaments may remain insoluble.

Informed by our ChIP-qPCR observations, we turned to ChIP-seq experiments to gain a broader understanding of SIR complex binding around the recombination event. We performed ChIP-seq during periods of active D-loop extension at 4 and 6 hour timepoints where we see the greatest increase in the ChIP-qPCR signal. These ChIP-seq signals were compared to a timepoint taken immediately before DSB induction (0 hour) where we do not observe enrichment of the SIR complex at donor or recipient loci. To assess the enrichment of the SIR complex accurately around the engineered recombination site, we used a modified S288C reference genome which includes the insertion of the engineered DSB and ectopic donor sites, as well as the knockouts of *HML*, *HMR*, *URA3*, *LEU2*, and *TRP1*, to represent the genome in a pre-DSB state. This more accurate representation of the genome, including engineered sites, allowed us to remove multi-mapped reads, and view uniquely aligned ones in the areas of interest, rather than having to blacklist non-unique sequences near the DSB site such as the *TEF* promoter (∼340 bp) and the 198 bp *TEF* terminator, which exist at multiple locations in this genome. We note that this processing filters out any reads mapping to the yeast telomere repeats, because such reads do not map uniquely. When evaluating our dataset, the first thing we note is the overall pattern of sharp peaks and valleys of signal (Fig 3). This pattern of signal is consistent with our method’s nucleosome footprinting effect and mirrors the pattern of enrichment seen in Thurtle and Rine, when performing Sir3 and Sir4 ChIP-seq using similar techniques to solubilize chromatin [72]. Consistent with our ChIP-qPCR data, we observe enrichment on the recipient strand upstream of the DSB at both 4 and 6 hour timepoints, as compared to the 0 hour control (Figs 3A and 3B; S3D Fig). While the overall increase in enrichment around the DSB is the greatest in the Sir2- and Sir3-TAP sample, each of the three subunits show enrichment in the area, suggesting the involvement of the full SIR2/3/4 complex, rather than the three playing independent roles. We suspect that the low enrichment of Sir4 in our experiment reflects our difficulty handling this subunit overall, as seen by our inability to visualize Sir4 via Western Blot.

**Fig 3.**
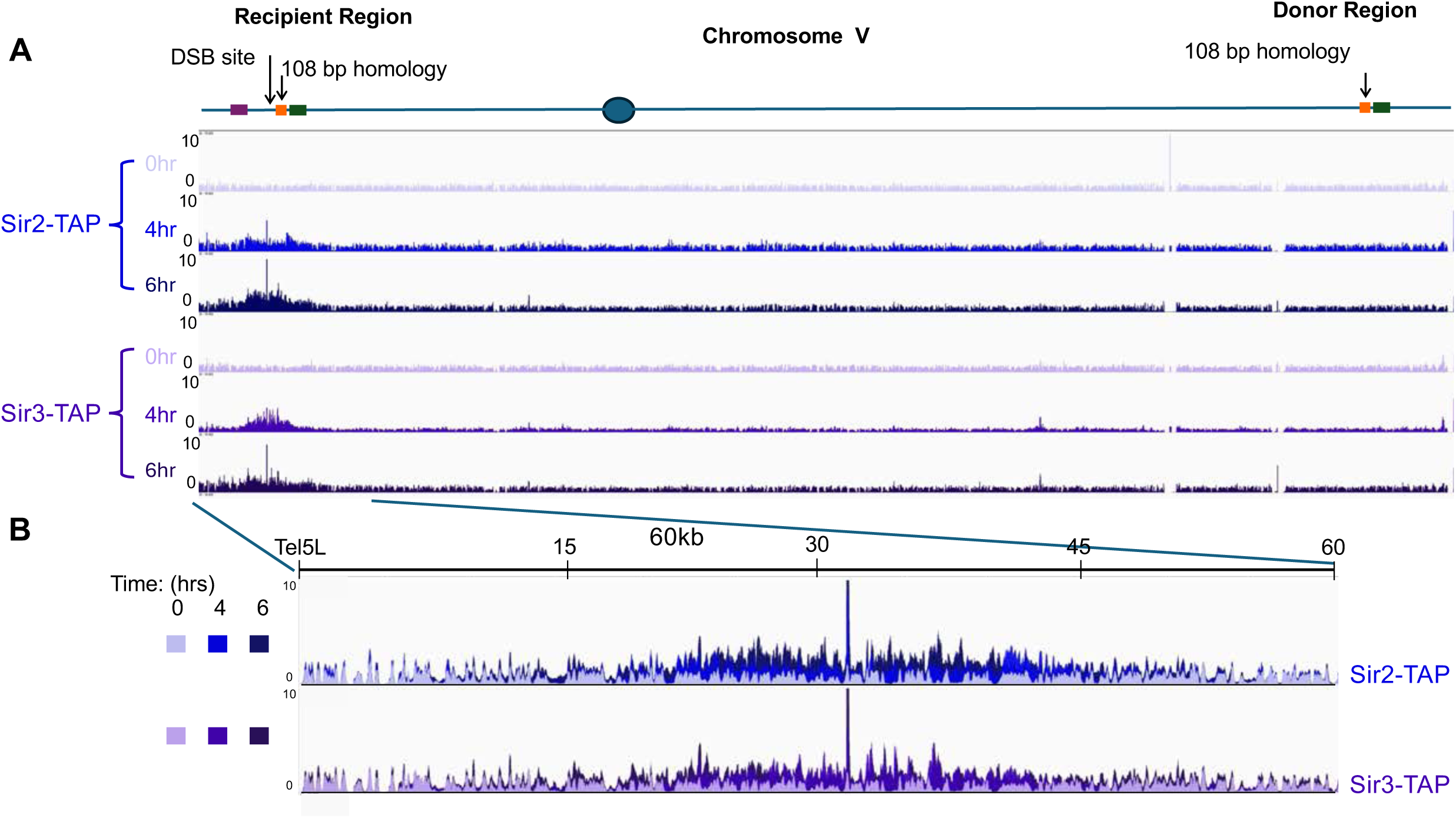
Sir2 and Sir3 enrichment at DSB during repair by BIR. (A) Chromosome-wide fold-enrichment signal for Chromosome V for Sir2-TAP, (Blue), Sir3-TAP (Indigo), at 0, 4, and 6 hrs post-DSB, as measured by ChlP-seq. Location of DSB indicated by schematic above data tracks. ChIP-seq signal is normalized by No-Tag controls (Materials and Methods). (B). Overlaid fold enrichment signals of Sir2-TAP and Sir3-TAP for 0, 4, and 6 hrs post-DSB induction. Window size is 60 kb from Tel5L. Height of all tracks scaled 0-10.

ChIP-seq reveals a symmetric SIR complex footprint that extends to roughly 15 kb from the DSB site; this binding pattern can be clearly observed when viewing the fold enrichment graphs of Chromosome V (Fig 3A, S3D Fig). This extensive binding domain explains, at least in part, the increased protein expression of Sir2 and Sir3, observed in Fig 1. Interestingly, in the ChIP-seq system, we do not observe significant binding at the donor locus. We believe this is due to the differences in sensitivity we observe between the two methods (see Discussion).

To characterize the timing and protein dependencies of SIR complex binding, we analyzed the recruitment of Sir2 in the absence of Rad51 and Rad52. BIR is significantly reduced in the absence of Rad51, and is ablated in the absence of Rad52, which is required for loading of Rad51 and its paralogs [73,74]. In *rad51Δ* strains, we observe binding of Sir2 immediately around the DSB, but the binding tract is reduced to roughly 7 kb (black arrow; Fig 4B) compared to 15 kb for *wild-type* (yellow arrow; Fig 4B). We also note that the residual binding tract appears asymmetric, with enrichment only on the centromeric side of the DSB. In *rad52Δ* strains, Sir2 enrichment is further reduced, displaying significant recruitment only in regions close to the DSB (Fig 4). Investigating the peak, which is independent of Rad51 and Rad52, we discover that it is located about 1 kb on the telomere side of the break, encompassing DNA from, and immediately distal to, the *TEF* terminator sequence (S5A Fig). We note that this is not likely to be an artifact of the genetic background, as non-uniquely mapped sequences were removed from analysis, and a similar peak is not observed at the *TEF* terminator which is located as part of the *KANMX* cassette used to remove the *LEU2* sequence in this genetic background (S5B Fig). Based on these observations, we conclude that there are two types of binding events occurring at the DSB; one which forms a small peak close to the DSB and occurs independent of repair pathway, and a broader domain of binding which is dependent on HR.

**Fig 4.**
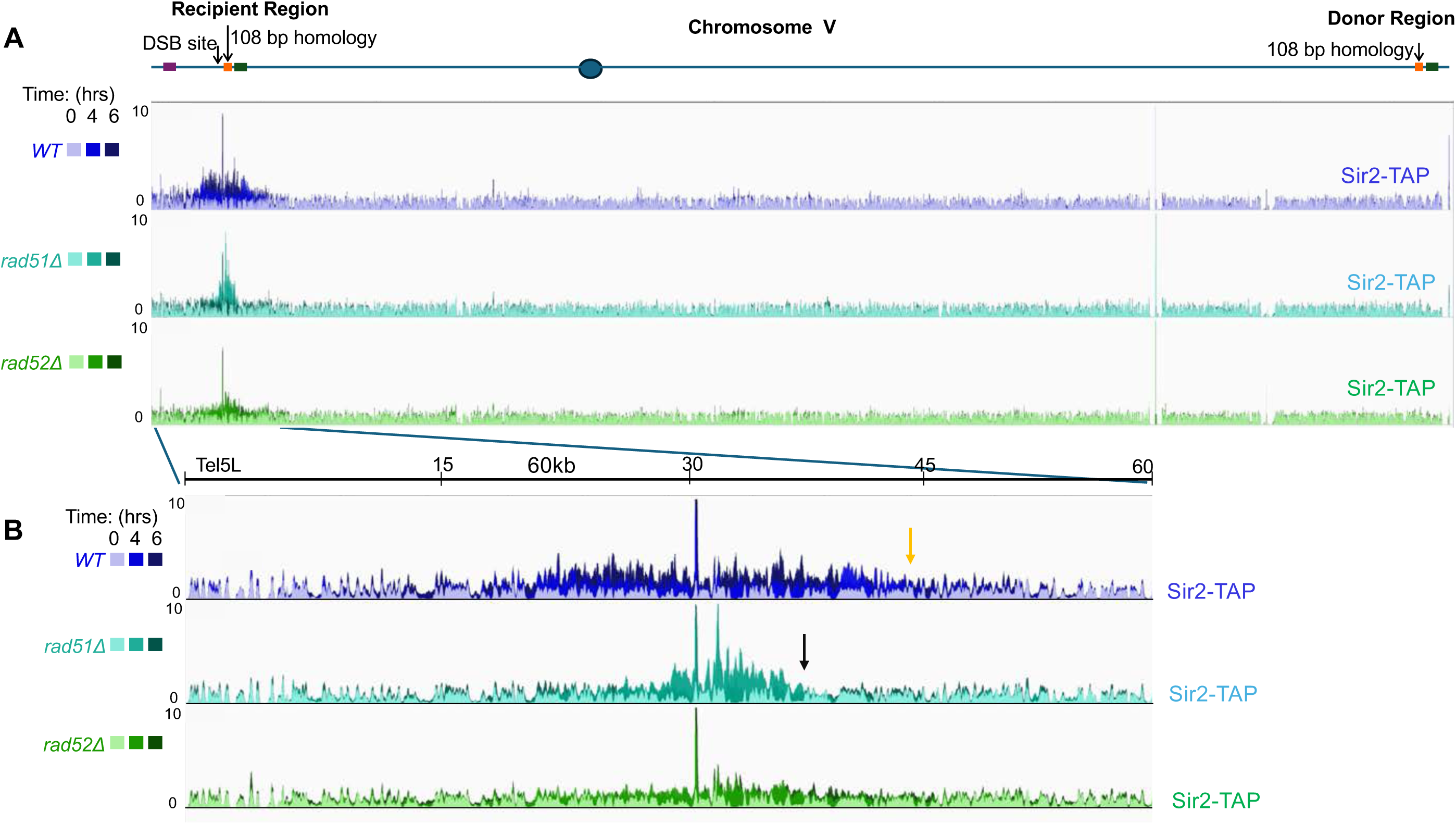
Sir2 enrichment at DSB in the presence and absence of proteins required for strand invasion. (A) Overlaid fold enrichment signals of Sir2-TAP for 0, 4, and 6 hrs post-DSB induction in *wild-type (WT), rad51Δ* and *rad52Δ* strains. Location of DSB indicated by schematic above data tracks. ChIP-seq signal is normalized by No-Tag controls. (B). Overlaid fold enrichment signals as in (A), with the window size 60 kb from Tel5L. Height of all tracks scaled 0-10. Arrows (gold for *WT*, black for *rad51Δ*) denote end of apparent enrichment track.

Another candidate for Sir2 recruitment is the mismatch recognition protein Msh2. Recruited early to DSBs, *msh2Δ* mutants show defects in the inheritance of silent chromatin regions [75,76].

Strikingly, in the absence of Msh2, we observe a substantial change in the binding of Sir2. Instead of forming relatively even distributions distal to the DSB, Sir2 appears to accumulate close to the DSB (Fig 5). In addition, we observe more Sir2 bound in the *msh2Δ* mutant. Thus, it appears that in this system, Msh2 seems to redistribute Sir2, removing or inhibiting its deposition from regions immediately about the DSB, and pushing enrichment farther upstream of the recombination intermediate.

**Fig 5.**
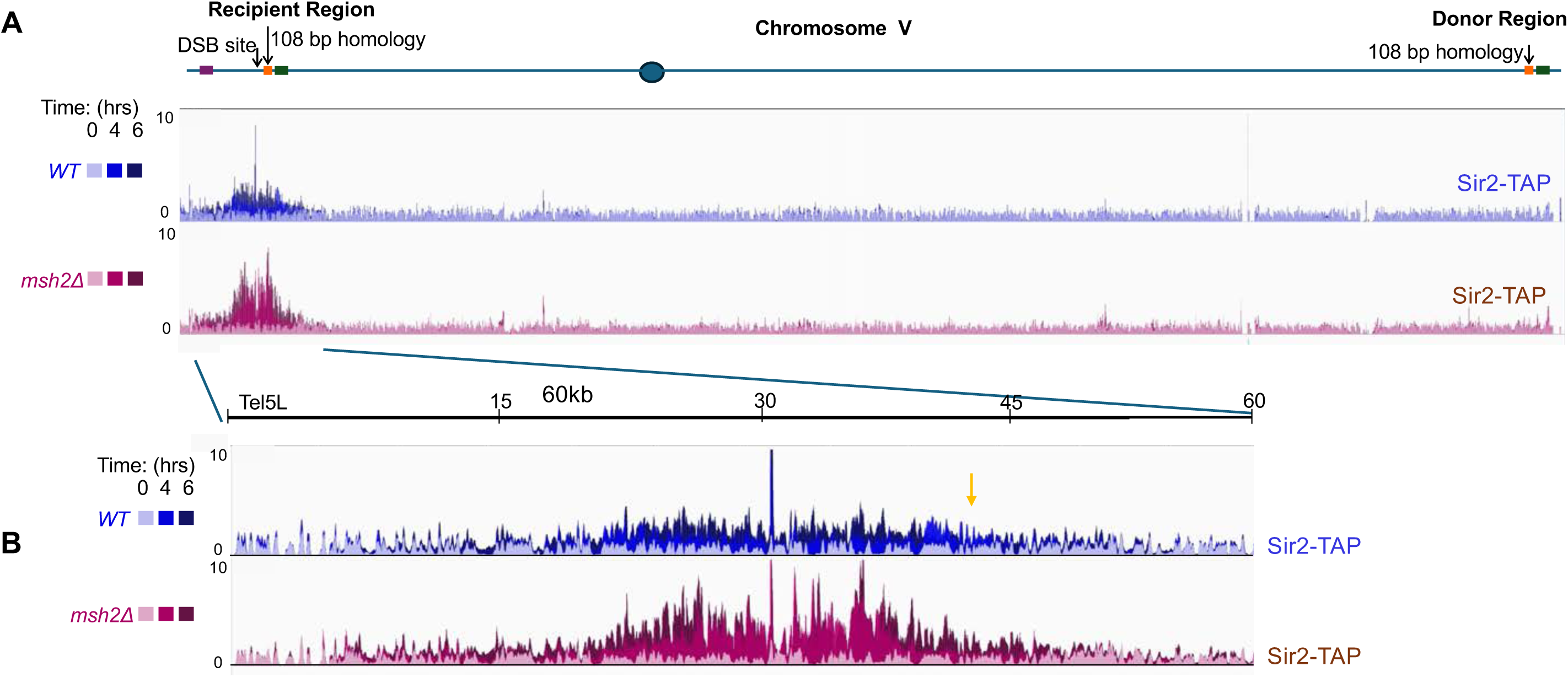
Sir2 enrichment at DSB in the presence and absence of Msh2. (A) Overlaid fold enrichment signals of Sir2-TAP for 0, 4, and 6 hrs post-DSB induction in *wild-type (WT)* and *msh2Δ* strains. Location of DSB indicated by schematic above data tracks. ChIP-seq signal is normalized by No-Tag controls. (B). Overlaid fold enrichment signals as in (A), with the window size 60 kb from Tel5L. Height of all tracks scaled 0-10.

This putative interaction between the SIR-bound domain and other HR related machinery is highlighted by an apparent defect in D-loop extension in Sir-TAP strains. Despite being functional for silencing [64–66] and haploproficient, Sir2-, Sir3- and Sir4-TAP strains all appear to display D-loop extension defects (Figs 2D and 2G; S3C Fig). We surmise that the TAP tag, when incorporated in the context of a SIR filament, inhibits the efficient interaction between the filament and recombination machinery.

Taken together, our results indicate upregulation of the SIR complex during BIR, allowing it to create a ∼15 kb domain upstream of the recipient and potentially donor strands. Given these observations, we hypothesized that the SIR complex might stabilize recombination intermediates during BIR to promote efficient extension, and that we might observe altered D-loop extension kinetics in the absence of the SIR complex. Thus, we set out to measure D-loop extension using the assay described in Fig 1A. For these experiments, we utilized one yeast strain which contained a homologous donor sequence and one which contained 9 mismatches across the 108 base pair donor homology (S1 Table). The strain containing mismatches in the donor is included to examine BIR under conditions where the stability of a strand invasion intermediate is compromised (Fig 1B). In the strain containing the homologous donor sequence, we do not observe any defect in D-loop extension in the absence of Sir2 or Sir4 (Figs 6B and 6D).

**Fig 6.**
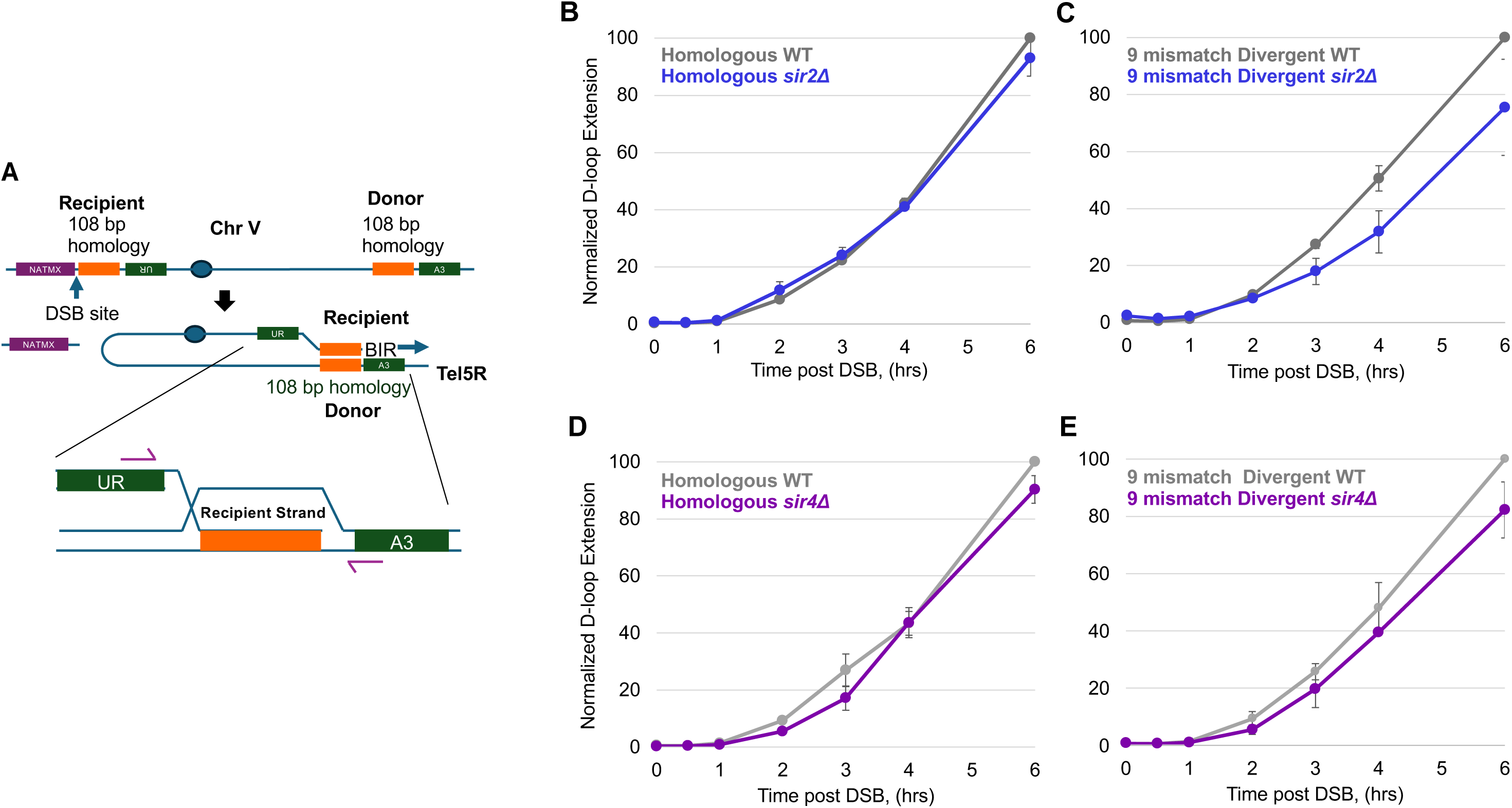
D-loop extension in BIR strains with and without Sir2 or Sir4. D-loop extension (A) for 0-6 hrs post-DSB induction for *wild-type* and *sir2Δ* and *sir4Δ* genotypes containing homologous donor (*WT* vs. *sir2Δ* (Panel B); *WT* vs *sir4Δ* (Panel D)) and a 9% divergent donor (*WT* vs. *sir2Δ* (Panel C); *WT* vs *sir4Δ* (Panel E)) sequences. D-loop extension values are normalized to *ZWF1* control locus, and then to the *WT* 6 hr data point. Error bars represent SEM.

However, in the strain containing a divergent donor sequence, we note slower D-loop extension in both *sir2Δ* and *sir4Δ* mutants (Figs 6C and 6E). This observation supports the hypothesis that the SIR complex plays a role in promoting efficient repair, especially in situations where a recombination intermediate would be unstable and prone to unwinding.

Next, we investigated the consequences of a slower BIR repair event. During the process of BIR, long stretches of single-stranded DNA are exposed on the recipient strand as the first strand is extended but the second strand has yet to be synthesized [23,24]. Single-stranded DNA is far more susceptible to damage than is double-stranded DNA, and the single-stranded nature of the recipient strand has been shown to lead to a higher rate of mutation on this strand as it is extended. Given these observations, we tested the rate of mutation during BIR in the presence and absence of the SIR proteins. To achieve this, we leveraged the selectable markers in the BIR recombination system [58] (Fig 7A). Upon DSB induction, BIR can repair one side of the DSB, which results in the formation of a functional *URA3* gene. Loss of the distal end of chromosome V is marked by the loss of the *NATMX* marker. Thus, if cells acquire a mutation in the latter half of the *URA3* gene as it is copied to the recipient during BIR, they would lose the *NATMX* marker, but fail to become Ura^+^. Inducing DSBs in mid-log cells and using a selection scheme to isolate these mutagenic BIR events, we observe an increase in the rate of such mutagenic BIR events in the *sir4* mutant (Fig 7E). Intriguingly, we observe the same trend in strains with homologous donor sequences, and those with 4 mismatches across the donor sequence.

**Fig 7.**
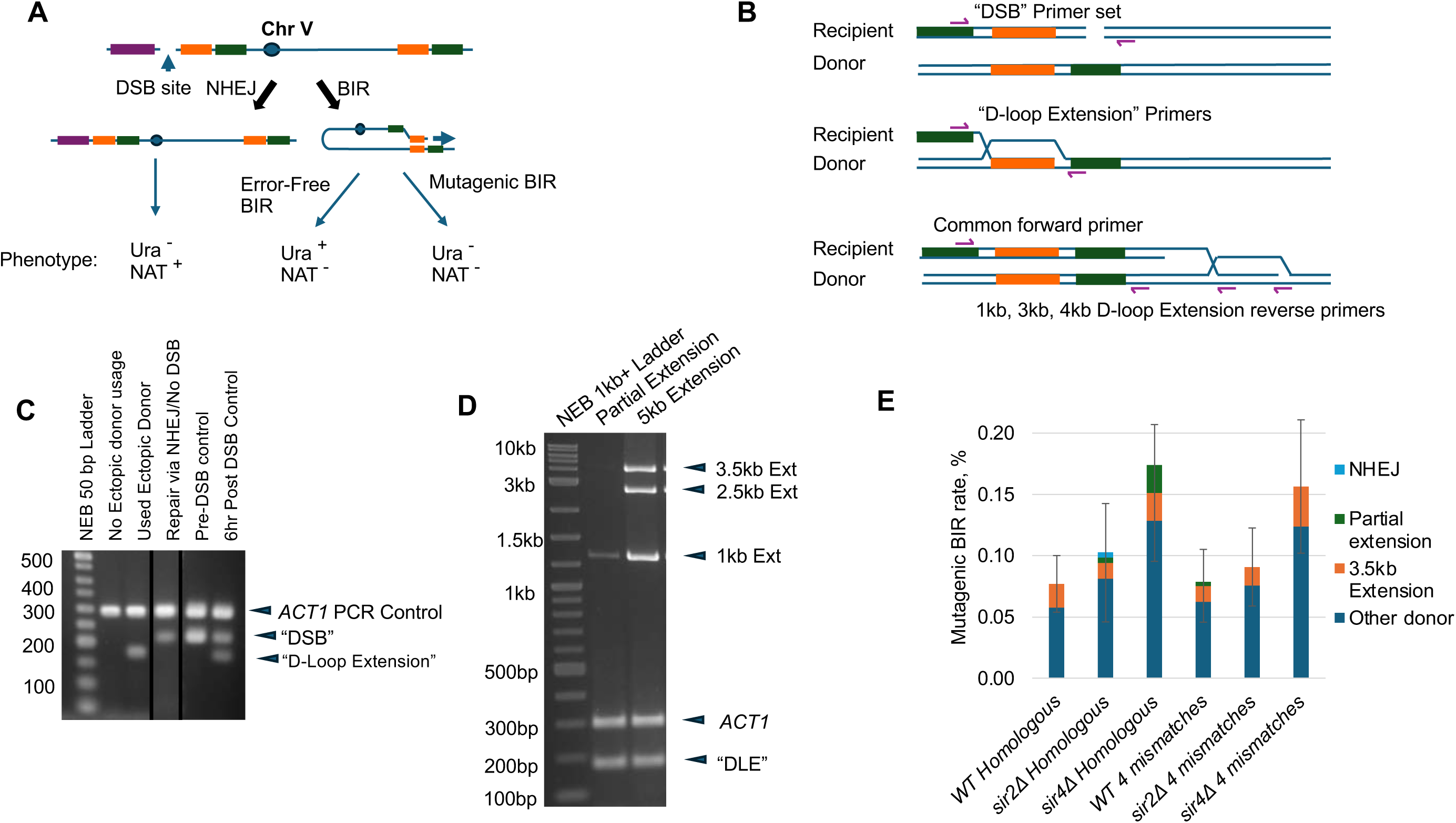
Quantification and characterization of mutagenic BIR events. (A) Schematic depicting selection for mutagenic BIR events. Upon DSB induction, cells. can repair via NHEJ, to remain resistant to nourseothricin, and Ura^-^. If cells repair the DSB via BIR in an error free manner, they become sensitive to nourseothricin, but Ura^+^. Repair by BIR in a mutagenic manner yields a nourseothricin^s^, Ura^-^ phenotype. (B) Diagram depicting PCR analysis of the mutagenic BIR survivor colonies selected for in a "DSB" primers produce amplicons in colonies which experienced no double strand break, or repair via "NHEJ. Invasion at the ectopic donor locus, and short-range extension yield the "D-loop extension” amplicon, while longer range extension to 1 kb, 2.5 kb, and 3.5 kb generate 1, 3, and 4 kb extension amplicons, respectively. (C) Examples of PCR results for colonies which performed NHEJ, BIR using the ectopic donor template, or mutagenic repair. (D) Examples of PCR results for colonies which performed BIR using the ectopic donor locus. (E) Mutagenic BIR rate for WT, *sir21′*, and *sir41′* genotypes with homologous and 3% divergent donor sequences. Survivor colonies were tested via PCR (Panels C and D), to rule out NHEJ, and determine ectopic donor usage, as well as level of extension using the ectopic donor. Error bars represent SEM. Strains analyzed (S1 Table) were yRA253 (homologous), EAY4697- 4699 (*sir2Δ*, homologous), EAY4706-4708 (*sir4Δ*, homologous), yAB280 (4 mismatches-3.7% divergent), EAY4703-4705 (*sir2Δ*, 4 mismatches), EAY4714-EAY4716 (*sir4Δ*, 4 mismatches).

To further investigate any differences between the *wild-type* and *sir* complex mutants in this context, we set out to test the donor template usage and sequence any mutations in the repaired chromosome immediately downstream of the donor locus (Fig 7B). To accomplish this, we tested 24 mutagenic BIR survivor colonies from *wild-type*, *sir2Δ* and *sir4Δ* backgrounds with either homologous or divergent donor sequences, using a PCR strategy shown graphically in Fig 7B. To first test whether the cells used the ectopic donor sequence and to confirm the absence of NHEJ, we performed PCR on each colony using a set of primers to detect either an uninduced DSB or an NHEJ product (“DSB” primers), primers which would amplify the chimeric molecule formed by D-loop extension at the donor sequence (“D-Loop Extension” primer set, also described in Fig 1A) and a loading control at an unrelated locus (*ACT1* on Chromosome VI; examples shown in Fig 7C). In accordance with our selection scheme, only one of the 144 survivor colonies displays a “DSB” amplicon. This confirms our ability to select against cells which did not induce a DSB or repaired said DSB via NHEJ. We suspect that the presence of this one colony indicates either cross contamination at the stage of picking survivor colonies from master plate, or mutagenesis of the *NATMX* gene, rendering the survivor colony sensitive to nourseothricin.

Surprisingly, most survivor colonies did not display any “D-loop extension” amplicon, indicating invasion and repair at a locus other than the 108 bp engineered ectopic donor sequence, or some other DNA repair strategy such as spontaneous telomere addition. This distribution does not appear to be different between *wild-type* and the *sir2Δ* or *sir4Δ* mutants. For each of the colonies which used the donor sequence as a template for BIR, we performed PCR to amplify the extended recipient molecule 1, 2.5, and 3.5 kb away from the donor template (Fig 7D). If a cell did initially invade at the donor locus, it is likely to extend at least 3.5 kb toward the telomere; in fact, only five of the 34 characterized survivor colonies failed to do so, extending 1 kb, but not further using the ectopic donor template. Sequencing these mutants reveals point mutations in the *URA3* gene in a proportion of the 5-FOA resistant colonies. These mutations were distributed equally across all genotypes (S8 Data, Underlying Data).

## Discussion

Here we characterize a new role for the SIR2/3/4 complex proteins in repair of DSBs by homologous recombination. To study these proteins in a direct manner, we utilized a BIR system to monitor the early stages of D-loop extension using qPCR and measure the localization of proteins to the donor and recipient strands upstream of the donor homology. In this system, we observe an upregulation of protein expression of Sir2 and Sir3 after DSB induction. Monitoring protein localization by ChIP-qPCR, we observe enrichment of the SIR complex at sites upstream of the region of homology on both the donor and recipient. We also note that these periods of enrichment closely correlate to times of active D-loop extension, as measured by the slope of the D-loop extension curves. Characterizing protein binding more extensively with ChIP-seq, we observe an extensive binding pattern in wild-type Sir2-, Sir3-, and Sir4- TAP strains, forming a 15 kb domain on each side of the DSB. While we focus on the centromeric side of the DSB, both sides of the DSB possess at least one donor locus elsewhere in the genome. Thus, the symmetric binding of Sir2, Sir3, and Sir4 is not surprising.

We hypothesize that a SIR complex of Sir2, Sir3 and Sir4 is recruited to a DSB. Such a process could occur through individual protein recruitment mechanisms; we note that Sir3 interacts with Sae2, which could explain its recruitment [52], and Sir4 can interact with the yKu70/80 complex independently of Sir3. Indeed, recruitment in such a manner might explain the early peak we observe very close to the DSB, which appears to be independent of the HR pathway. However, we also note that the Ku complex is required for the establishment of silent chromatin domains bound by the Sir2/3/4 complex [75]. The extensive binding tract of each subunit, extending up to 15 kb from the DSB site, suggests to us the formation of a silent chromatin domain, rather than individual subunit recruitment in a 1:1 manner by Sae2 and the yKu complexes. Further, if recruitment of individual subunits to the DSB by Sae2 and the Ku complex were to solely explain this phenomenon, we would not expect recruitment of the 15 kb domain to be dependent upon HR. Further evidence against a 1:1 binding mode comes from previous work by the Gasser lab, who observe Ku protein enrichment dwindling by the time the bulk of the Sir3 and Sir4 enrichment begins at a DSB [54]. Lastly, our ChIP-qPCR experiments, which appear to be more sensitive than our ChIP-seq studies, show localization of Sir2, Sir3, and Sir4 upstream of the donor template during repair by BIR, where we would not expect any yKu70/80 binding, or any action by Sae2. Thus, while we certainly do not rule out the role of Sae2/Sir3 interactions or yKu70/80/Sir2/4 in this system, we do not think that this explanation is sufficient to explain our observations.

Taken together, our observations suggest a model where the SIR complex displays two binding modes. First, it is recruited to the immediate vicinity of the DSBs, independently of DSB repair. Once repair has started, a more extensive silent chromatin domain is established upstream of the extending repair event on the recipient chromosome. This domain spreads to the donor sequence, perhaps due to its proximity (Fig 8). What proteins are involved in the establishment of this SIR domain remains an open question: candidates such as yKu70/80 are attractive due to their presence at DSBs and their roles in establishing silent chromatin elsewhere [47,54]. It is also possible that the SIR complex is recruited via the Sir3-Sae2 interaction [52]. Further, the relationship between the SIR complex and the DNA damage-associated γH2A histone phosphorylation remains an intriguing question. While regions bound by the SIR complex are constitutively marked by γH2A phosphorylation, mutation of H2A serine 129 does not abrogate SIR binding to those regions. Rather, mutation of *sir3* results in the loss of γH2A phosphorylation at these silent loci [77]. We hope that future work may help to answer these questions, as well as clarify the roles of individual SIR subunits vs the complex as a whole in the DSB repair process.

**Fig 8.**
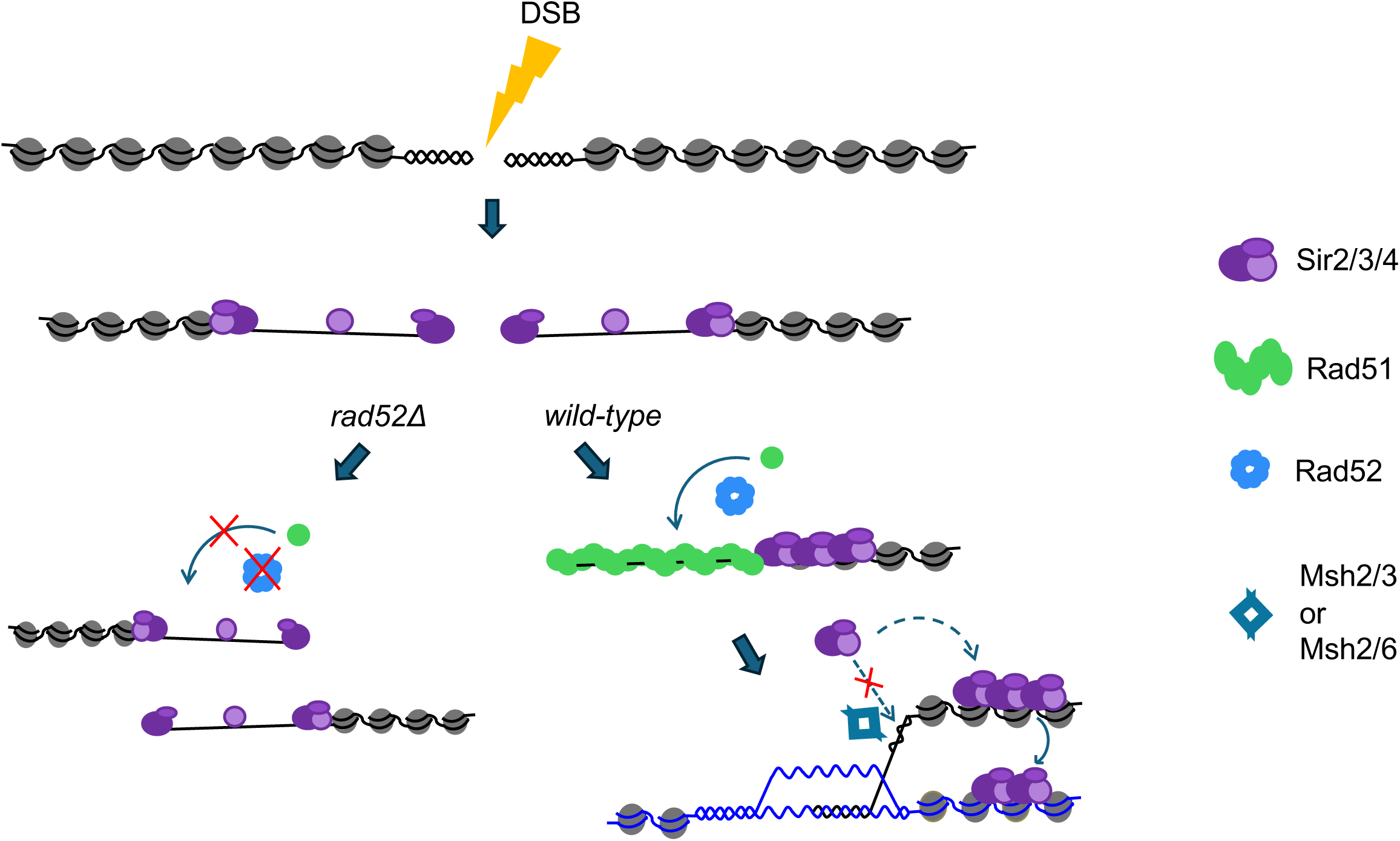
**Model, depicting an initiating DSB, which is bound by Sir2 and Sir4, possibly via yKu70/80, and Sir3, potentially through known Sir3/Sae2 interactions**. In the absence of Rad52, strand invasion does not occur, and thus no major change is observed. In HR competent cells, Rad52 loads Rad51 to form a nucleoprotein filament, which invades at a donor locus, leading to increased SIR2/3/4 complex recruitment. These SIR complexes are either relocalized or inhibited from being deposited immediately behind the extending D-loop by the action of Msh2. SIR domain formation on the recipient locus upstream of the break spreads to the donor locus, stabilizing the recombination intermediate, leading to more rapid D-loop extension.

This two-part binding model reflects our observations of Sir2 binding in the absence of proteins necessary for strand invasion, such as Rad51 and Rad52. In such mutants, we note that while recruitment of Sir2 very close to the DSB remains, the extent of the Sir2 binding tract is roughly halved when compared to *wild-type* at 4 hours. At the 6 hour timepoint, binding is further diminished, whereas wild-type binding remains significant. These observations support a model where initial recruitment near the DSB is independent of repair, but that during the recombination process, a larger SIR bound domain is established farther from the repairing recombination intermediate.

A surprising observation is the change in binding pattern of Sir2 in the absence of the mismatch repair factor Msh2. We note a significant increase in Sir2 localization in *msh2Δ* mutants, and a shift in its pattern of localization. Rather than forming a relatively even distribution ∼2-15 kb distal to the DSB at early timepoints, we observe significant enrichment 1-15 kb from the DSB, which only seems to strengthen between the 4 and 6 hour timepoints. This indicates that either Msh2 plays some sort of inhibitory role in the deposition of Sir2, especially close to the DSB site, or that Msh2 relocates Sir2, and possibly the other SIR complex members, to form a filament farther from the DSB. Hints of a relationship between DNA mismatch repair components and silent chromatin have been observed previously; Msh2 was shown to be required for silencing at *HMR*, and its loss was shown to cause Sir2 torelocate from the SIR complex-bound *HMR/HML* and telomeres to the rDNA locus [75]. Liu et al. [75] observe that the absence of the mismatch repair proteins Msh2, Mlh1 or Pms1 causes a loss of yKu70 at *HMR*, leading to failure to recruit and establish a silent chromatin domain. However, how these factors might interact in the presence of a DSB remains unclear and difficult to predict. This interplay between repair factors and the SIR complex is highlighted by the D-loop extension defect observed in the Sir-TAP strains, despite their competence for silencing.

At the donor locus, we observe enrichment in our ChIP-qPCR experiments, but we do not see such enrichment reproducibly in our ChIP-seq experiments. We hypothesize that this is due to the significantly lower sensitivity in the ChIP-seq experiments compared to their ChIP-qPCR counterparts. One potential explanation for this difference is that while qPCR should be able to detect any precipitated DNA fragments larger than 50-80 bp, the library preparation steps of our ChIP-seq experiments size selects for 120-300 bp fragments. Thus, any larger DNA fragments which proved refractory to nuclease digestion due to SIR complex binding would not be visible to our ChIP-seq. Second, ChIP-qPCR experiments should be able to detect single-stranded DNA relatively efficiently, whereas ChIP-seq experiments would not. Lastly, the data processing and analysis of ChIP-qPCR and ChIP-seq experiments differ, making it difficult to perform head-to-head comparisons of the two techniques. Thus, we believe that there is enrichment of the SIR complex at the donor locus, but that this enrichment falls below the limit of detection in the ChIP-seq experiments.

Our results suggest that once established, this SIR bound silent chromatin domain promotes efficient repair. In instances where repair is highly inefficient, such as BIR events with non-homologous donor sequences, we observe defects in the speed of D-loop extension in the absence of Sir2 and Sir4.

These observations are supported by our observation that, *sir4Δ*, and possibly *sir2Δ* genotypes appear to leave the cell vulnerable to mutation accumulation during BIR, as indicated by an increase in the rate of mutagenic BIR in the survival assay.

The SIR complex might stabilize recombination intermediates through several mechanisms. First, it could establish transcriptionally silent regions on recipient and donor sequences which would suppress transcription mediated destabilization. This hypothesis is supported by recent observations by the Piazza lab, who demonstrate D-loop dissolution by transcription through donor sequences [78] . Second, SIR complex bound regions of the genome are known to interact with one another [69]; such tethering might serve to stabilize the extending D-loop or promote re-invasion at the same locus in the event of D-loop dissolution. Third, SIR bound domains are known to normally localize to the nuclear periphery [34,43,45,46,79–81], but Sir2 and Sir3 have each been shown to relocate away from the nuclear periphery to DSBs in the nuclear lumen upon DSB induction [54,55]. In such a scheme the SIR complex binds luminal DSBs and may assist in the localization of the DSB to the nuclear periphery as it would to any other silent chromatin domain [34,35,43,45,70], perhaps participating in the SUMO dependent relocalization of slow to repair DSBs to the periphery [82,83]. Once there, Sir4’s tethering to the nuclear periphery via Esc1 [43,45,46,79,80] might have a similarly stabilizing effect on recombination intermediates. This hypothesis is especially attractive, given the larger effect of the *sir4Δ* mutation on mutagenic BIR in the survival assay.

## Materials and Methods

### Media

*S. cerevisiae* yeast strains used in this study were grown at 30°C in either yeast extract peptone-dextrose (YPD) or synthetic complete media supplemented with 2% lactate (pH 5.5), or 2% galactose [84]. When required, geneticin (Invitrogen, San Diego), nourseothricin (Werner BioAgents, Germany) or Hygromycin B (Invitrogen) were added to media at recommended concentrations [85].

### Strain Construction

All strains, plasmids, and oligonucleotides used in this study are described in S1, S2 and S3 Tables, respectively. For *sir2Δ* and *sir4Δ* genotypes: strains containing homologous (yRA253) or 9% divergent (yRA275) donor strains were transformed using the standard lithium acetate transformation technique [86] with plasmids containing Cas9 and guide RNAs targeting the Sir2 ORF or Sir4 ORF, respectively (S2 Table), along with repair templates consisting of regions upstream and downstream to the ORF fused together, without the Sir2 or Sir4 ORF. Plasmid transformation was selected by plating cells on minimal media with monosodium glutamate [87] and nourseothricin but lacking ammonium sulfate and uracil. Plasmids were then removed by plating colonies onto minimal media containing 5-FOA (5-Fluoroorotic acid, Thermo Fisher Scientific; [84]). Colonies were then struck to singles on YPD + nourseothricin media, and *sir2Δ* and *sir4Δ* genotypes were confirmed by PCR with the oligos listed in S3 Table. For these and all strains generated in this study, oligos used to confirm genotypes lie outside those used to generate amplicons transformed into each strain.

For *msh2Δ*, *rad51Δ* and *rad52Δ* genotypes: Integration vectors were digested with restriction enzymes described in S2 Table and transformed into appropriate genotypes using the standard lithium acetate transformation method [86], selecting for integrants with minimal media lacking leucine.

Colonies were then struck to singles on minimal media lacking leucine, and genotypes were tested using the oligos outlined in S3 Table. For *msh2Δ* transformant genotyping, where amplicon size between *MSH2* and *msh2Δ::LEU2* were very similar, amplicons were digested with *Bgl*II.

For Sir2-, Sir3-, and Sir4-TAP genotypes: Strains from the yeast TAP collection [61,62] contain each gene, C-terminally tagged with a glycine linker, followed by the *HIS3MX* selectable marker. These strains were transformed with pAG34 (S2 Table) digested with *Fsp*I, and integrants were selected on YPD+Hygromycin to replace the *HIS3MX* selectable marker with *HYGMX*. The *SirX-TAP-HYGMX* fragments amplified with oligos described in S3 Table, and transformed into EAY4460, and selected on YPD + hygromycin media to generate *SirX-TAP-HYGMX* in the BIR background. Transformant genotypes were confirmed by PCR with oligos described in S3 Table.

### BIR Survival Assay

BIR survival assays were performed as described Anand et al [58,59]. Strain yRA253 was struck from glycerol stock onto YPD + nourseothricin media. After two days, ∼2 mm in diameter colonies were resuspended in 1 mL sterile water. 10 µL of this resuspension was then diluted 100-fold, into 990 µL of sterile water. 50 µL of this dilution was plated onto YPD and YP+2% galactose plates. After two days, YPD plates were counted after two days, to ascertain number of cells per colony, and YP-2% galactose plates were replica plated onto minimal Ura dropout media and grown for two more days. Ura^+^ colonies were counted, and BIR frequency was calculated as number of Ura^+^ colonies divided by number of colonies on YPD.

### Testing Haploproficiency of Sir-TAP strains

Single colonies of FY91 (*MAT*alpha) were mated to single colonies of BY4741 (*MAT*a, *wild-type*), EAY4056 (*MATa, sir2Δ*), EAY5412 (*MATa, SIR2-TAP*), EAY5413 (*MATa, SIR3-TAP*) and EAY5414 (*MATa, SIR4-TAP*) (S1 Table). Mated cells were grown overnight (∼17 hrs) on YPD plates. Each mating mixture was resuspended into 100 µL of sterile dH2O after which ten-fold serial dilutions were made. 5 µL of the 10^-2^, 10^-3^, 10^-4^, and 10^-5^ dilutions were spotted onto YPD and minimal Uracil, Adenine dropout plates. Plates were grown at 30°C and photographed after 36 hours on YPD and 48 hours on minimal Uracil, Adenine dropout plates. Three independent matings were performed for each genotype.

#### BIR time courses

Cells were struck from glycerol stocks to single colonies on YPD + nourseothricin plates and grown at 30°C for two days. Single colonies were then inoculated into 5 mL YP + 2% lactate + nourseothricin liquid cultures. After ∼24 hours of growth, 4.5-9 Optical Densities600 (ODs) of cells were inoculated into 250 mL (for BIR qPCR time courses) or 400 mL (for Western blots, ChIP time courses, or ssDNA degradation qPCR experiments) of YP + 2 % lactate media, and grown overnight. To induce DSBs, galactose was added to a final concentration of 2%. 10 (for BIR qPCR time courses), 20 (for ssDNA degradation experiments and western blot time courses) or 30 (for ChIP experiments) ODs of cells were collected at timepoints indicated. Cells for 0 hour timepoints were collected immediately prior to galactose addition. To collect each timepoint, cells were collected and immediately placed on ice, pelleted at 3000 RPM in swinging bucket centrifuge, and washed with 10 mL of chilled TBS (20 mM Tris-HCL pH 7.5, 150 mM NaCl). Cells pellets were resuspended in 1 mL chilled TBS, transferred to microcentrifuge tubes, re-pelleted, flash frozen in liquid nitrogen, and stored at -80°C. For ChIP experiments, after cell collection, cells were cross-linked at room temperature for 15 minutes with formaldehyde to a final concentration of 1%. Formaldehyde was quenched at room temperature for 5 minutes with 2 M glycine to a final concentration of 300 mM. Cells were then placed on ice and washed as described above.

### Western blotting

Flash frozen cell pellets were resuspended on ice in 100 µL lysis buffer (150 mM NaCl, 25 mM Tris-HCl pH 8, 1 mM EDTA, 10 mM β-mercaptoethanol, 1x HALT protease inhibitor (Thermo Fisher Scientific), to which 200 mg acid washed glass beads (500 mM, Sigma) were added. Cells were vortexed at 4°C for 10 minutes. 5x SDS-PAGE loading buffer (0.25 M Tris-HCL pH 7, 0.5 M DTT, 10 % SDS, 50 % glycerol, 0.5 % bromophenol blue) was added to a to 1X final concentration and incubated at 95°C for 5 minutes. 12-20 µL of sample was loaded on 1.5 mM thick 6-15% SDS-PAGE gels, then transferred to 0.45 micron PVDF membranes (Thermo Fisher Scientific). Transfer was confirmed via staining with ponceau and blocked with 5% milk in 1X TBST (20 mM Tris-HCL pH 7.5, 150 mM NaCl, 0.1% Tween- 20). Primary antibodies were diluted appropriately (1:1,000 for anti-CBP (Sigma-Aldrich), 1:10,000 for GAPDH (Sigma-Aldrich) and incubated at 4°C overnight. After three washes with 1X TBST, membranes were incubated with appropriately (1:20,000 to 1:40,000 dilution of secondary antibody in block solution), diluted HRP-conjugated secondary antibodies (goat anti-Rabbit-HRP, Thermo Fisher Scientific, and rabbit anti-mouse-HRP, Sigma-Aldrich), washed 3 times in 1X TBST again, and imaged using a LI-COR Odyssey M Imager (Lincoln, NE). Blots were quantified relative to GAPDH loading control using Fiji [88]. For analysis by Western blot, three independent transformants were used to generate three separate time courses on separate days. Western blots were then performed on these independent time courses on separate days.

### qPCR analysis

DNA was extracted using standard Phenol::Chloroform methods [89]. Briefly, cell pellets were resuspended in 100 µL of lysis buffer (10 mM Tris-HCL pH 8, 1 mM EDTA, 100 mM NaCl, 1% SDS, 2% Triton x-100), and lysed by vortexing for 10 minutes at 4°C with 0.2 g acid washed glass beads. Lysis buffer was diluted with 200 µL of 1x TE (10 mM Tris pH 8, 1 mM EDTA), and RNA was digested with 2.5 µL of RNAse A at 37°C for 45 minutes. DNA was extracted into equal volumes of phenol:chloroform:isoamyl alcohol (25:24:1), then into chloroform. To the aqueous phase, 5 M NaCl was added to a final concentration of 200 mM, 1 µL of glycoblue (Thermo Fisher Scientific) was added to visualize the DNA pellet, and DNA was precipitated in 2.5 volumes of 100% ethanol. DNA pellets were washed in 70% ethanol, resuspended in 20 µL of ultrapure water, and quantified via the Qubit broad range dsDNA quantification kit (Thermo Fisher Scientific). 1-2.5 ng of DNA was used for downstream qPCR analysis. qPCR reactions were conducted 10 µL reactions in triplicate, with the following conditions, with Phusion polymerase (New England Biolabs) and SYBR green (Thermo Fisher Scientific): Initial denaturation, 98°C for 30 seconds, followed by 40 cycles of denaturation at 98°C for 10 seconds, annealing at 65°C for 25 seconds, and extension at 72°C for 15 seconds (Roche LightCycler 480 II). For qPCR analysis, D-loop extension Cp values were normalized by *ZWF1* controls. For D-loop extension time courses, three independent transformants of the *sir2Δ* and *sir4Δ* transformants were each grown on separate days. Each replicate was compared to a wild-type replicate (S1 Table) [58].

### ExoVII treatment of DNA samples from BIR time courses

DNA samples were prepared for qPCR analysis, but volumes of cells and reagents were doubled, to obtain enough DNA for analysis. 100 ng of DNA was digested in a 10 µL reaction containing 1 unit of ExoVII (New England Biolabs) in (10 mM Tris-HCL pH 8, 50 mM KCl) at 37 °C for 30 minutes, then then heat inactivated at 95 °C for 15 minutes. Samples were then diluted 1/10, 1 µL of which was used in each qPCR reaction. Untreated controls were treated identically, except ExoVII was omitted. D-loop extension was calculated as described in the qPCR section of the Materials and Methods. In this experiment, non-specific digestion of dsDNA by ExoVII is controlled for by normalization of the D-loop extension signal by the double-stranded *ZWF1* signal (outlined in the previous section).

### ChIP experiments

Cell pellets were resuspended in 400 µL of lysis buffer (50 mM HEPES-KOH, pH 7.5, 1 mM EDTA, 150 mM NaCl, 0.1% sodium deoxycholate, 0.1% SDS, 1% Triton X-100, 1X HALT protease inhibitor), and transferred to 2 mL screw-top tubes containing 0.5 g acid washed glass beads. Lysis buffer was added, and tubes were closed in a way to minimize air bubbles. Cells were then lysed in a bead beater for a total of (24) minutes of bead beating, at intervals of 3 minute on, with 5 minutes rests on ice. Lysate was transferred to a 15 mL conical tube, and chromatin was pelleted by centrifugation at 4,000 rpm for 5 minutes in a Beckman J6 centrifuge. The resultant chromatin pellet was resuspended in 1 mL of lysis buffer, and magnesium was added to a final concentration of 1.5 mM to facilitate nuclease digestion.

Lysate was removed from ice, and 4 units of Pierce Universal Nuclease (Thermo Fisher Scientific) was added to each tube and incubated at room temperature for 15 minutes. Nuclease digestion was halted by the addition of EDTA to a final concentration of 10 mM, and tubes were returned to ice. An equal volume (1 mL) of Dilution buffer (50 mM HEPES-KOH, pH 7.5, 100 mM NaCl, 0.1% sodium deoxycholate, 0.05% SDS, 1% Triton X-100, 1X HALT protease inhibitor) was added to the sheared lysate, which was incubated on a rotisserie at 4°C for 10 minutes. Lysate was then spun at 3000 RPM in a microcentrifuge for 5 minutes to remove debris. 2% input samples were removed at this time, and the rest of the cleared lysate was removed to a tube containing 0.6 mg (estimated 3-6 µg Rabbit IgG) Rabbit IgG conjugated Dynabeads M270 epoxy (Thermo Fisher Scientific), which had been pre-washed in lysis+dilution buffer 1:1. Samples were incubated with rotation overnight at 4C. Beads were then washed 4x with ChIP wash buffer (100 mM Tris-HCL pH 7.5, 0.5 M NaCl, 2 mM EDTA, 1% IPEGAL-CA 630, 1% sodium deoxycholate, 1X HALT protease inhibitor), then twice with 1X TE. Samples were then resuspended in 1X TE, (2% input samples were brought up to 100 µL with TE), and RNA was digested by addition of 2µL of 20 mg/mL RNAse A, and incubation at 37°C for 20 minutes. To each tube was added SDS to a final concentration of 1%, and 2 µL of proteinase K, 10 mg/mL. Samples were then uncross-linked by incubation at 65°C overnight. DNA was then extracted by phenol::chloroform in the same manner as the qPCR samples previously described. For ChIP-qPCR and ChIP-seq experiments, 2-3 independent transformants were utilized, with the exception of the *wild-type* (yRA253) and *msh2Δ* no-tag strains (yAB256) obtained from the Haber lab [58]. Experiments for each independent transformant of a particular genotype were performed on separate days.

### ChIP-seq library preparation

For ChIP-seq experiments, BIR time courses were performed on separate days using a minimum of two independent transformants per genotype. Library prep was accomplished using an Illumina Tru-seq strategy (Illumina, San Diego, CA), and transformants were processed on separate days. First, DNA was end-repaired and 5’ phosphorylated by incubation for 15 minutes at 25°C, then 30 minutes at 37°C in a reaction with the following final concentrations: 1X T4 DNA ligase buffer, 0.33 µM dNTPs, 0.035 units/µL Large Klenow fragment (New England Biolabs), 0.1 units/µL T4 polymerase (New England Biolabs), 0.35 units/µL T4 polynucleotide kinase (New England Biolabs). End-repaired, 5’ phosphorylated DNA fragments were then purified using AMPure SPRI magnetic beads (Beckman Coulter, Indianapolis IN) at a ratio of 2:1. A-tailing was accomplished by incubation for 30 minutes at 37°C in a reaction with the following final concentrations: 1X CutSmart (New England Biolabs), 0.1 mM dATP, 0.3 units/µL Small Klenow fragment. This reaction was then heat inactivated by incubation at 75°C for 20 minutes. 0.12 pmol of New England Biolabs indexed adaptors were added to each sample, and adaptors were then ligated by addition of the following components, to a final concentration: 8.4% w/v PEG 8000, 1 mM ATP, 1X CutSmart, 20 units/µL T4 DNA ligase (New England Biolabs). Samples were incubated for 30 minutes at room temperature. DNA libraries were then purified again with AMPure SPRI beads at a 2:1 ratio. Libraries were amplified using 15 cycles of PCR with NEB universal primer, and Q5 DNA polymerase (New England Biolabs) using the manufacturer’s specifications.

Libraries were then purified from adaptor dimers by TBE-PAGE followed by gel extraction, selecting amplicons from ∼250-500 bp. Briefly, crushed gel pieces were incubated in 3 volumes (600 µL) of gel elution buffer (10 mM Tris-HCL pH 8, 0.5 mM ammonium acetate, 10 mM magnesium acetate, 1 mM EDTA, 0.1% SDS), and incubated overnight at 37°C with rotation. Supernatant was removed, and a second extraction was performed with 1.5 volumes (300 µL) of gel elution buffer, for 4 hours. Eluate was distributed into two tubes, and sample was ethanol precipitated with 1 µL glycoblue, 16.66 µL of NaCl, and 2.5 volumes (usually about 1 mL). DNA pellets were washed in 75% ethanol, dried, and resuspended in ultrapure water. DNA concentration was achieved by means of a Qubit high-sensitivity DNA quantitation kit. For ChIP-qPCR experiments, PCR was conducted using the same conditions as those described in the “qPCR analysis” section. Paired-end libraries were sequenced on an Element Aviti DNA sequencer (Element Biosciences, San Diego, CA), sequencing 80 bp of each insert. The raw sequence data are deposited in the National Centre for Biotechnology Information Sequence Read Archive under accession number PRJNA1262862.

### ChIP-seq analysis

In this study, we analyze SIR complex binding at an engineered recombination site. To accurately observe these phenomena, we first sequenced areas of difference from the S288C reference genome by Sanger sequencing (Cornell BioResource Center) and edited the S288C R64 reference genome (https://www.ncbi.nlm.nih.gov/datasets/genome/GCF_000146045.2/) to reflect the genetic background (yRA253, see S1 Table). This was accomplished by means of the ReForm tool [90] which generated GFF annotation files and FASTA sequences for the new reference genome. Sequenced library quality was confirmed by FastQC. For each genotype, replicates were pooled and analyzed as one dataset. Libraries were aligned to this reference genome using Bowtie2, using “local”, “sensitive” presets. PCR duplicates were then removed with Samtools. Reads with a MAPQ score of less than 2 were then removed from the dataset, which had the effect of removing low quality reads, those that mapped with equal scores to multiple genomic locations (“true multimappers”), and those with greater than or equal to four mismatches at high certainty bases [91]. After these filtering steps, each replicate contained a minimum of 500,000 reads. The average number of reads per replicate at each time point was 12 million. As a control, all of the no-tag replicates of a particular genotype (*wild-type*, *msh2Δ*, *rad51Δ*, *rad52Δ*) at a particular timepoint were combined to form a “master no-tag” control for a particular genotype, at a particular timepoint. We refrain from combining genotypes any further due to potentially different DSB resection kinetics. Normalization factors for each pair-wise comparison of SirX-TAP to Master No-tag datasets were obtained by NCIS using Scriptmanager [92,93]. To generate tracks for peak calling/visualization, control tracks were generated from the Master No-tag BAM files using MACS3 subcommands to replicate the function of MACS3 callpeaks function, as is recommended in the MACS3 documentation when utilizing MACS3 for DNAse-seq experiments [94,95]. When constructing the background (lambda) tracks, we utilized a fragment size (d) equal to the average fragment size of the experiments to which the background track was to be compared, as determined by the MACS3 predicted function in paired-end mode. Slocal and llocal values were set to 500 bp and 5 kb, respectively, and the global background was omitted from the generation of the background track lambda, because of the observed loss of signal around the DSB as a result of DNA end resection. For each pair-wise comparison, Sir2X-TAP samples were scaled to their respective background tracks using the NCIS derived scaling factors, by scaling the larger dataset down to the smaller dataset, using the MACS3 BDGopt function.

These scaled bedgraphs were then used to generate fold-enrichment tracks using the MACS3 bdgcmp function.

### Mutagenic BIR survival assay

Cells were struck from glycerol stocks to single colonies on YPD + nourseothricin plates and grown at 30°C for two days. Single colonies were then inoculated into 5 mL YP + 2% lactate + nourseothricin liquid cultures and grown for ∼24 hours. 2 mL YP + 2% lactate cultures were inoculated with 0.03 ODs of culture and grown to mid-log overnight (∼16 hours). Mid-log cultures were then diluted (1:50 for synthetic minimal media + 5-FOA +2% galactose plates, 1:2,500 for YPD plates), and 100 µL of each dilution was plated onto YP-2% galactose plates to induce a DSB, and YPD plates to ascertain the number of cells per culture. After incubation at 30°C for two days, colonies on YPD plates were counted, and this number was used to determine the concentration of cells in each culture. After three days, DSB survivors on the 5-FOA + 2% galactose plates were replica plated onto YPD + nourseothricin plates, and then onto YPD plates. These replica plates were then counted the following day. YPD + nourseothricin colony numbers were subtracted from the replica plated YPD colony numbers, to obtain the number of mutagenic BIR (5-FOA^r^, Nat^s^) events. The number of mutagenic BIR events was then scaled by the dilution factor, and the number of colonies per culture to obtain a mutagenic BIR rate per culture. 30-33 individual cultures were analyzed in this manner, spread across 6 experimental days. Three independent transformants were used for *sir2Δ* and *sir4Δ* genotypes, and these were compared to *wild-type* strains obtained from the Haber lab (S1 Table; Fig 7 legend) [58].

### Mutagenic BIR colony survivor analysis

DNA was extracted from cultures grown from mutagenic BIR survivor colonies using a standard Phenol::Chloroform method [89]. First, we performed PCR using three PCR primer sets: as a positive control PCR to the actin locus, “*ACT1*” with AO4777 and 4778 (expected amplicon size 292 bp). To test whether the repair event utilized the 108 bp ectopic donor sequence, amplified with the D-loop extension primer set (AO4771 and 4772, expected amplicon size 173 bp). Lastly, to test whether NHEJ was performed, the “DSB” primer set was utilized (expected to produce an amplicon if either a) DSBs were not induced, or b) the DSB was repaired by NHEJ, (AO4764 and 4765, expected size 220 bp. As DNA controls, 0 and 6 hr DNA samples from the population repair assays were utilized to confirm PCR was possible at all three loci. PCR was performed using 40 cycles of 63°C annealing temp, and 30 seconds of 72°C extension. A second round of PCRs were performed on colonies displaying possibly positive “DLE” amplicons to confirm their donor template usage. For the second round, PCR was performed using the *ACT1* amplicon as a positive control, the “DLE” amplicon to confirm donor locus usage, and three amplicons which would be expected to amplify if the colony in question continued to extend the D-loop 1 kb, 2.5 kb, and 3.5 kb from the donor locus. Based on the presence or absence of the “DSB”, “DLE”, “1 kb extension”, “2.5 kb extension”, and “3.5 kb extension” amplicons, colonies were characterized as having performed NHEJ (“*ACT1*” + “DSB” amplification), BIR using another donor template (“*ACT1*” amplification alone), Partial extension (“*ACT1*”, “DLE” and potential “1 kb extension”, “2.5 kb extension” amplification), or extension to 3.5 kb (“*ACT1*”, “DLE” and 1 to 3.5 kb extension primer sets). For those colonies which displayed D-loop extension to at least 1 kb, amplicons were sequenced by Oxford Nanopore sequencing by Plasmidsaurus (South San Francisco, CA).

## Acknowledgements

We thank William Lai and Amy Lyndaker for advice and providing Aviti sequencing for ChIPseq experiments, Philip Versluis and Abdullah Ozer for discussion regarding ChIP- based techniques sequence analysis and normalization, and Wolf Heyer and the Heyer lab for stimulating discussion regarding D-loop extension and its detection. We thank Maria Bravo Nunez for comments on the manuscript, Jim Haber and Frank Pugh for generously providing strains, and Jeff Pleiss for generously providing access to his qPCR machines.

## Supplemental Figure, Supplemental Table, and Underlying Data Captions

**Supplemental Fig 1.**
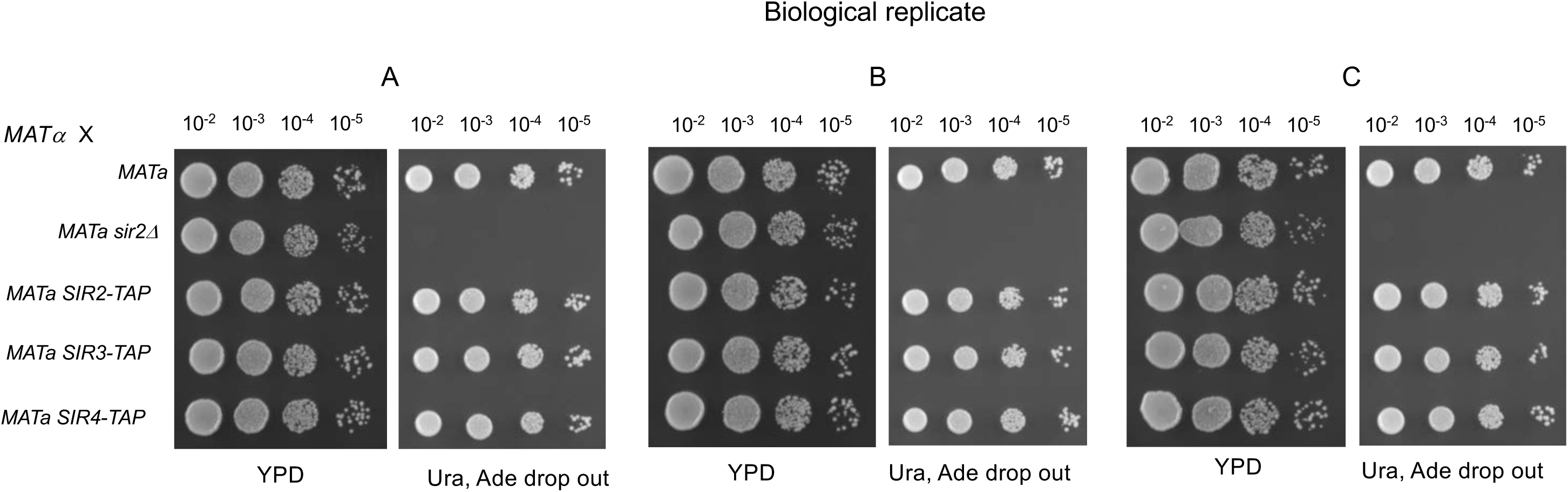
Mating type assay used to assess haploproficiency of Sir2- Sir3- and Sir4-TAP strains. Mating assay for *wild-type, sir2Δ, Sir2-TAP, Sir3-TAP*, and *Sir4-TAP* (all *MATa*) strains with *MATα* mating type tester (Materials and Methods). Three independent biological replicates were assayed (Replicates A, B, and C)

**Supplemental Fig 2.**
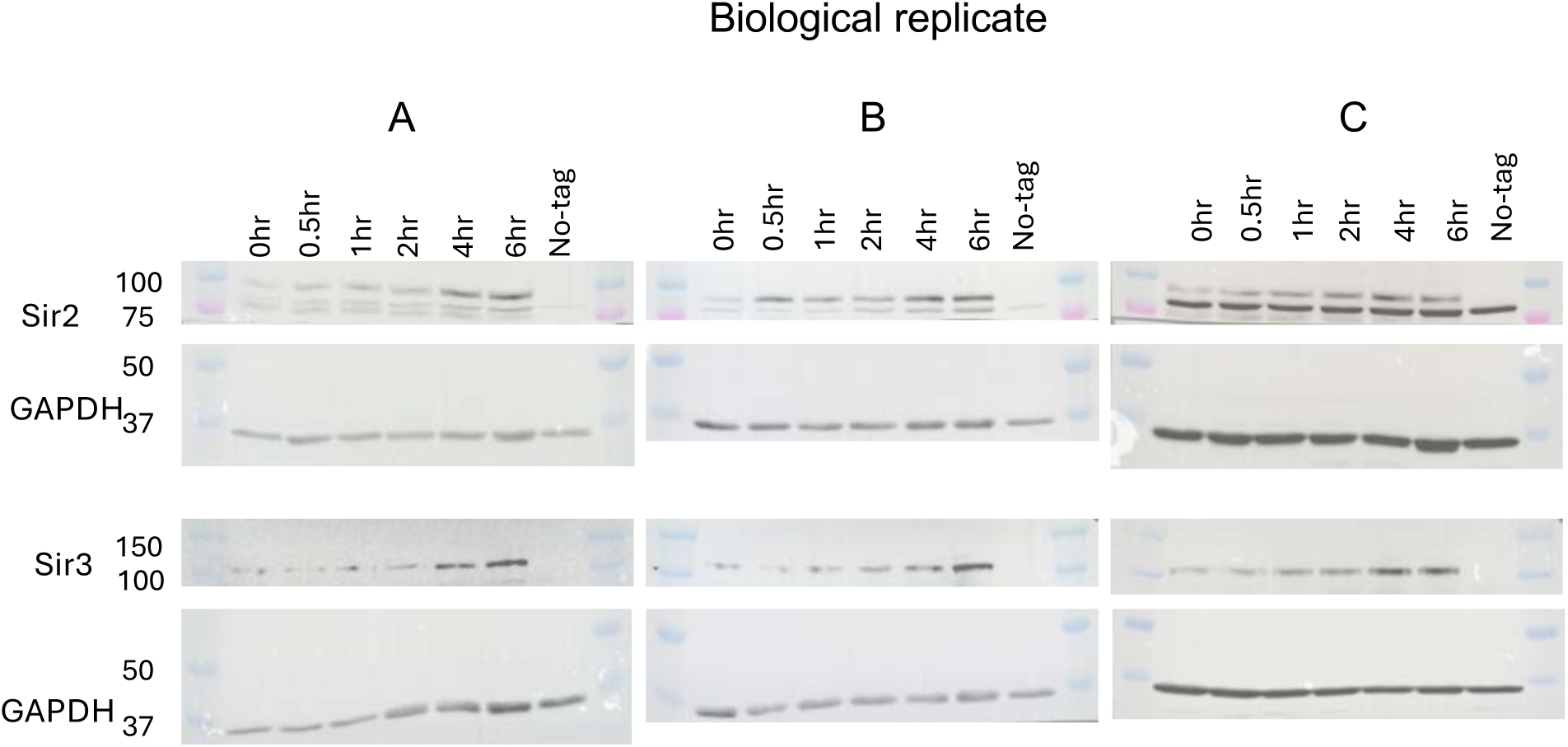
Western blots measuring Sir2 and Sir3 proteins levels during BIR. Independent transformants of Sir2- and Sir3-TAP were assayed in independent experiments. Western blots were conducted on samples 0-6 hrs post DSB induction, and Sir2- and Sir3-TAP protein signals were normalized by GAPDH loading controls.

**Supplemental Fig 3.**
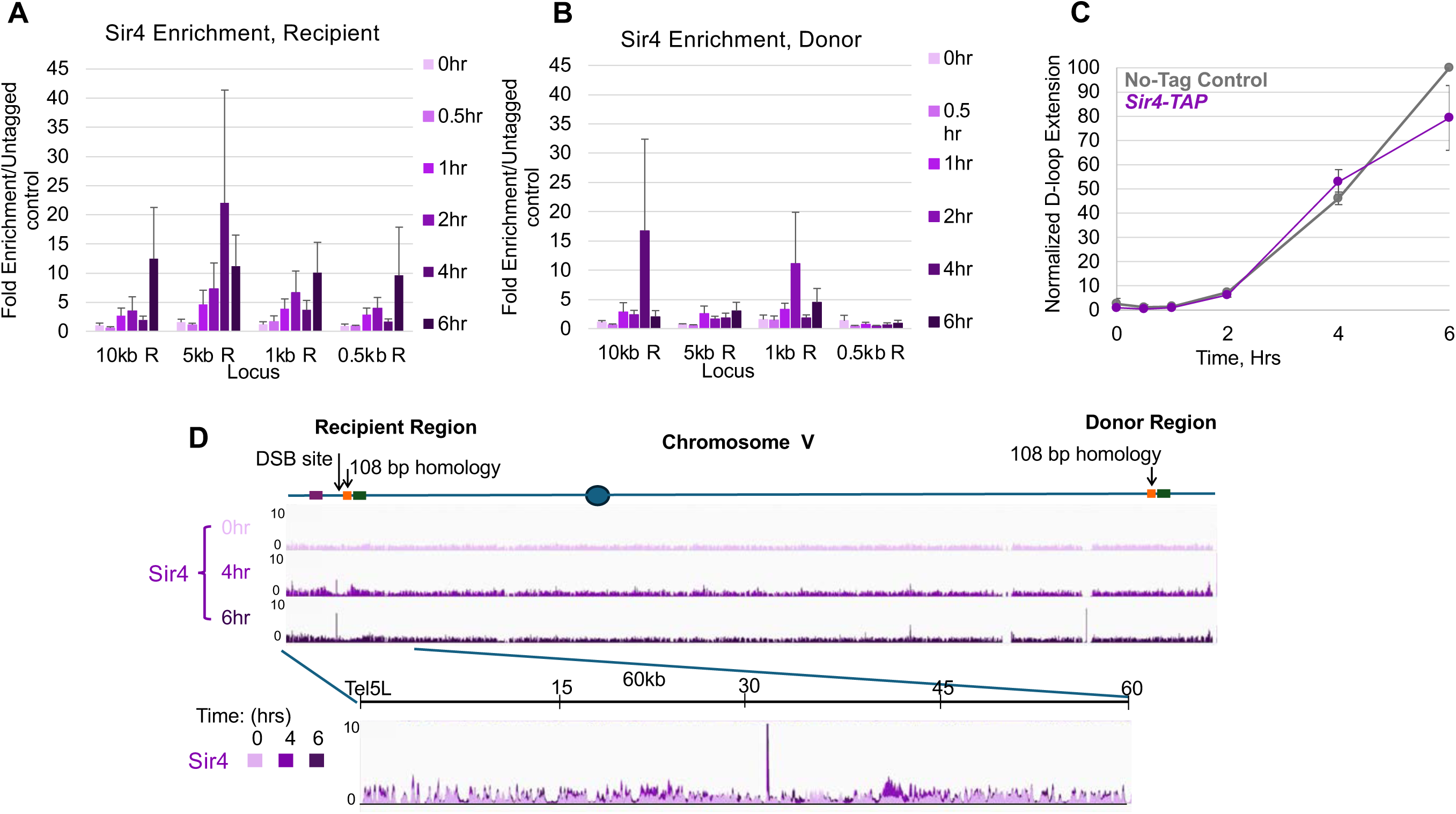
Enrichment of Sir4 at the DSB during BIR. Average fold enrichment over No-Tag control strains for Sir4-TAP at recipient (A) and donor (B) loci. at 0.5, 1, 5, and 10 kb upstream of the recipient and donor loci. (C) D-loop extension data corresponding to the time courses in (A) and (B). For the ChIP and D-loop extension experiments, averages are the result of three independent transformants, assayed on separate days. Error bars represent SEM. (D) Overlaid fold enrichment signal of Sir4 for 0, 4, and 6 hrs post-DSB induction. Window size is 60 kb from Tel5L. Height of all tracks scaled 0-10.

**Supplemental Fig 4.**
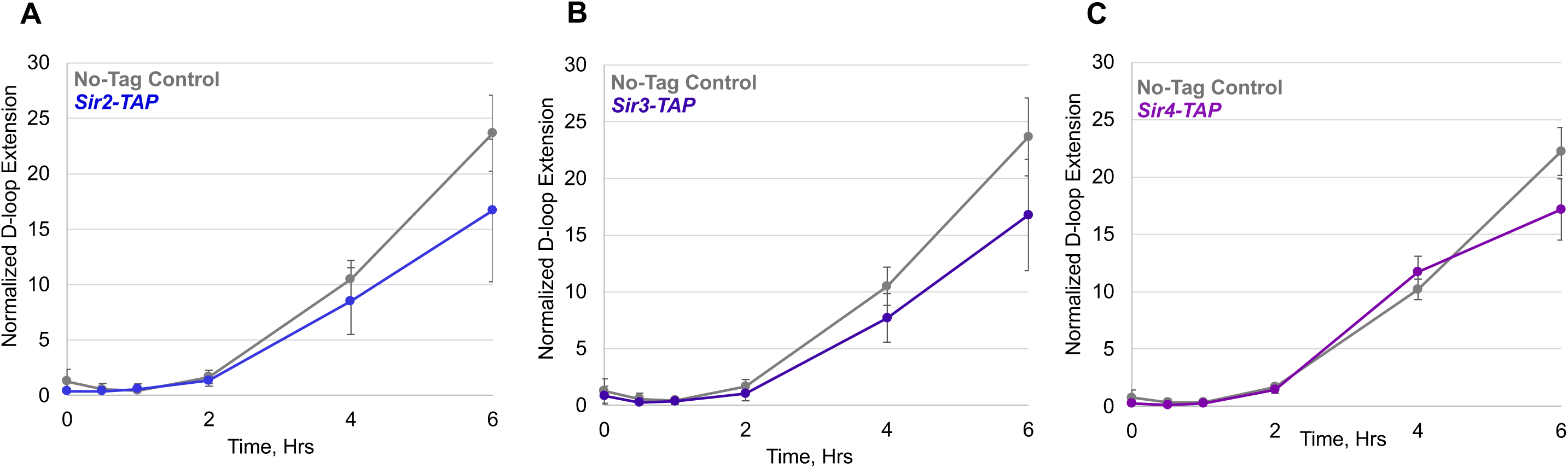
Raw D-loop extension during BIR. D-loop extension for Sir2-TAP and No-Tag control (A), Sir3-TAP and No-Tag control (B), and Sir4-TAP and No-Tag control (C), without normalization by No-Tag controls (Materials and Methods). No-Tag control averages represent replicates performed in parallel to the SirX-TAP strains to which they are compared. Error bars represent SEM.

**Supplemental Fig 5.**
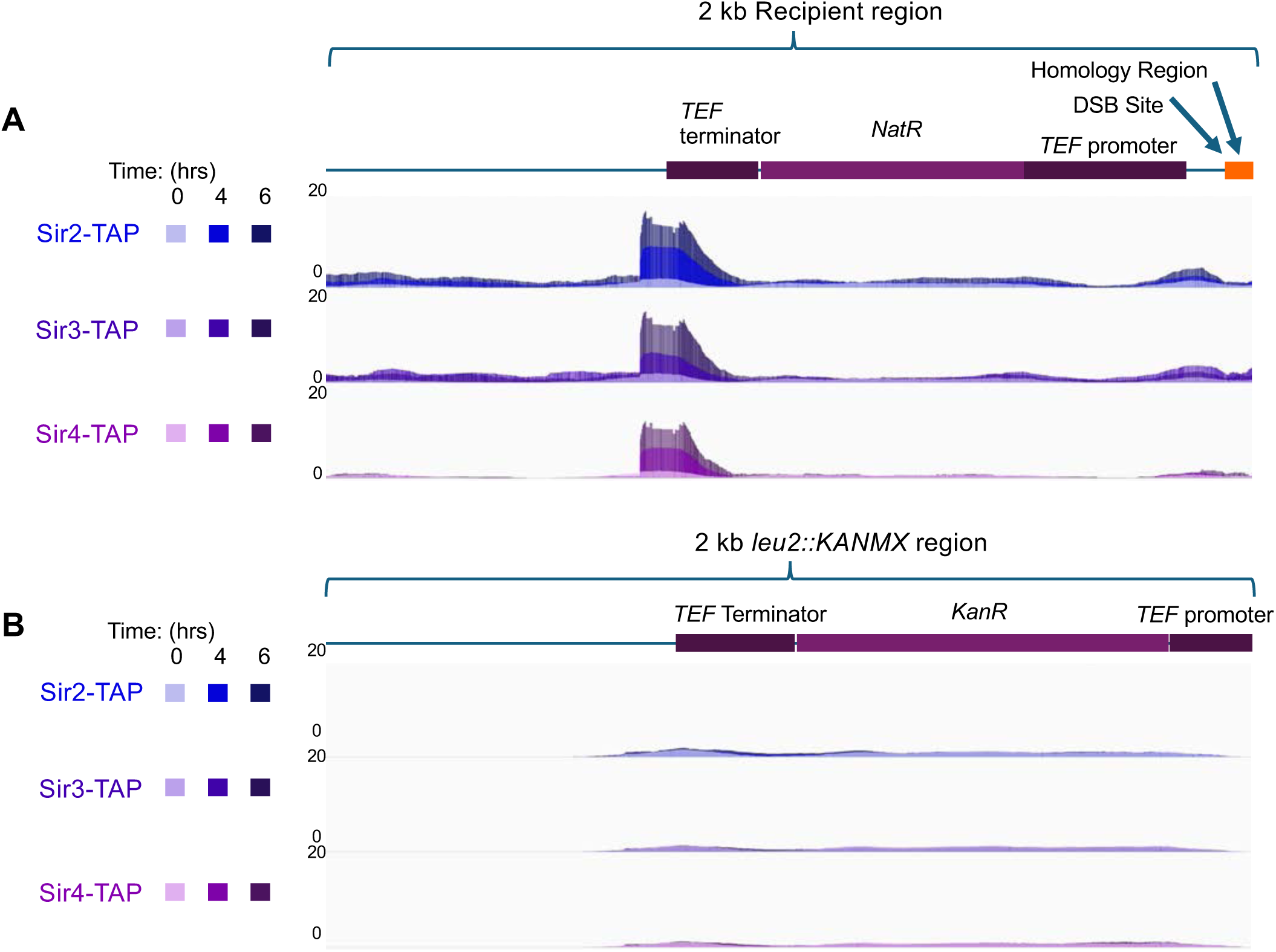
SIR complex enrichment near DSB site vs Chr III *KANMX* region. (A) 2 kb window displaying Sir2, Sir3, and Sir4 enrichment near the *TEF* terminator in the *NATMX* cassette near the DSB site on Chr. V. (B) Equivalently sized window showing enrichment of Sir2, Sir3, and Sir4 at the *KANMX* cassette, used to replace *LEU2* on Chr. III.

**S1 Table.**
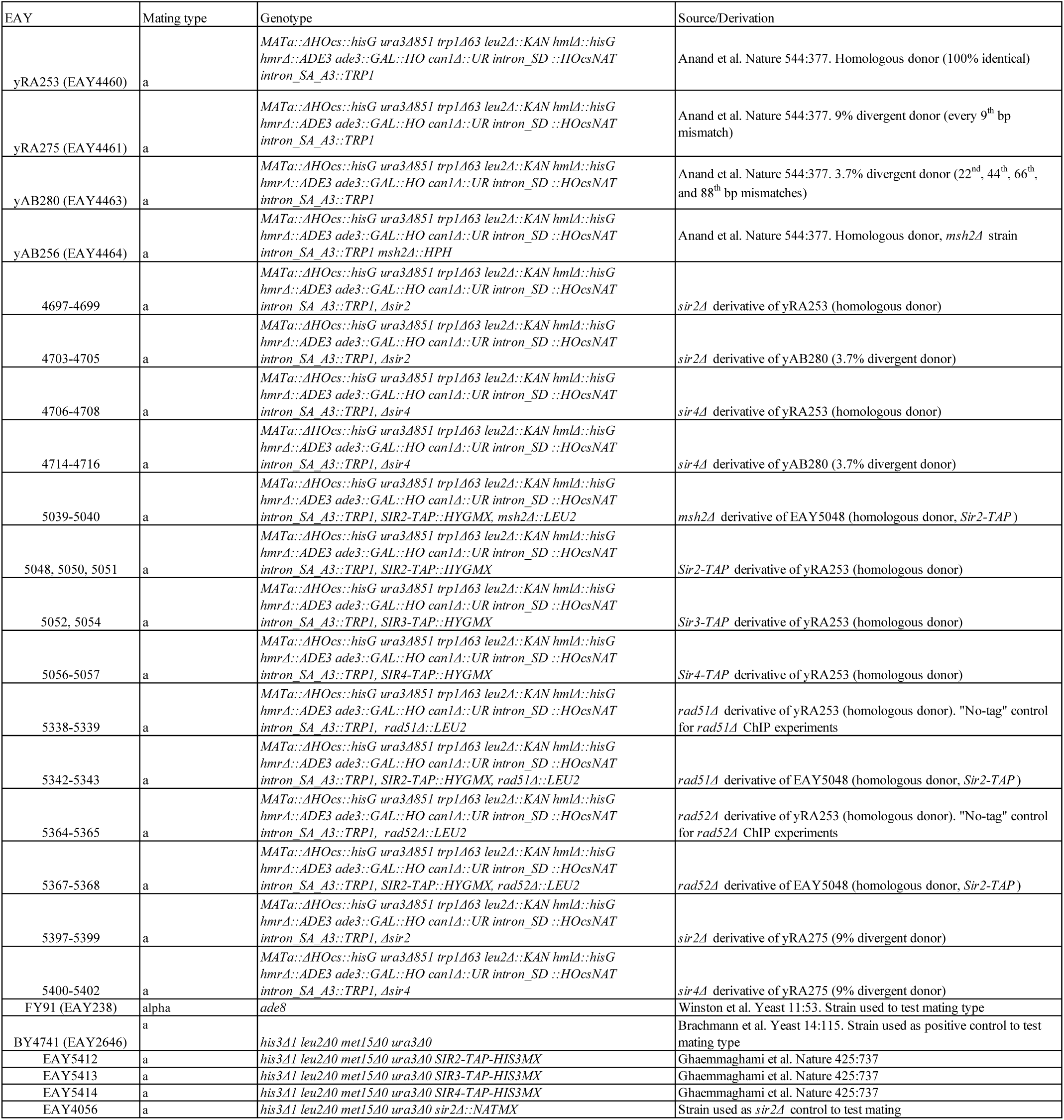
Strains used in this study.

**S2 Table.** Plasmids used in this study.

| Name | Description |
| --- | --- |
| pEAM334 | 2 micron yeast origin, *ampR*, *pUC origin* . Contains *URA3* , *CAS9* , guide RNA sequence targeting *SIR2* ORF. |
| pEAM335 | 2 micron yeast origin, *ampR*, *pUC origin*. Contains *URA3* , *CAS9* , guide RNA sequence targeting a second location in the *SIR2* ORF. |
| pEAM336 | 2 micron yeast origin, *ampR*, *pUC origin*. containing *URA3* , *CAS9* , guide RNA sequence targeting *SIR4* ORF. |
| pAM50 (pEAO178) | *ampR*, *pUC origin* integration vector. Digest with *Bam* HI and *Pst* I, to release a 5.6 kb *rad51Δ::LEU2* fragment for integration. Obtained from James Haber laboratory. |
| pAG34 (pEAO162) | Hygromycin B resistance vector (Euroscarf) |
| pBRDHSLEU2 (pEAO74) | *ampR*, *pUC origin* integration vector. Digest with *Nhe* I and *Sph* I to release *rad52Δ::LEU2* fragment for integration. Contains upstream region of *RAD52* , followed by ~ 3 kb fragment containing *LEU2* under its native promoter, followed by downstream region of *RAD52* . Long read sequencing reveals plasmid to be a dimer, which does not affect its function. Obtained from Dennis Livingston. |
| pEAI110 | *ampR*, *pUC origin* integration vector. Digest with *Aat* II and *Pvu* II to release a 4.4 kb *msh2Δ::LEU2* fragment for integration. |

**S3 Table.**
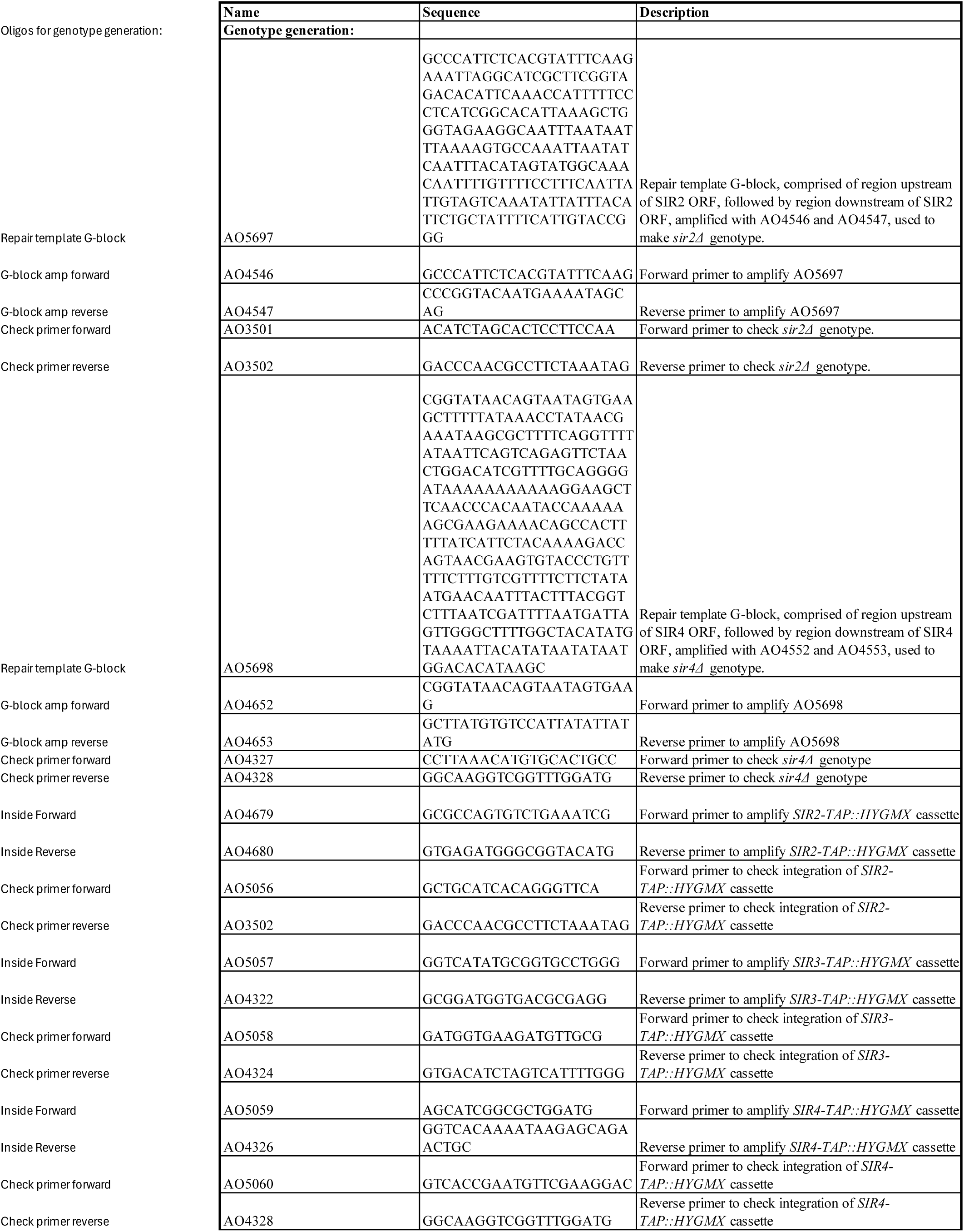
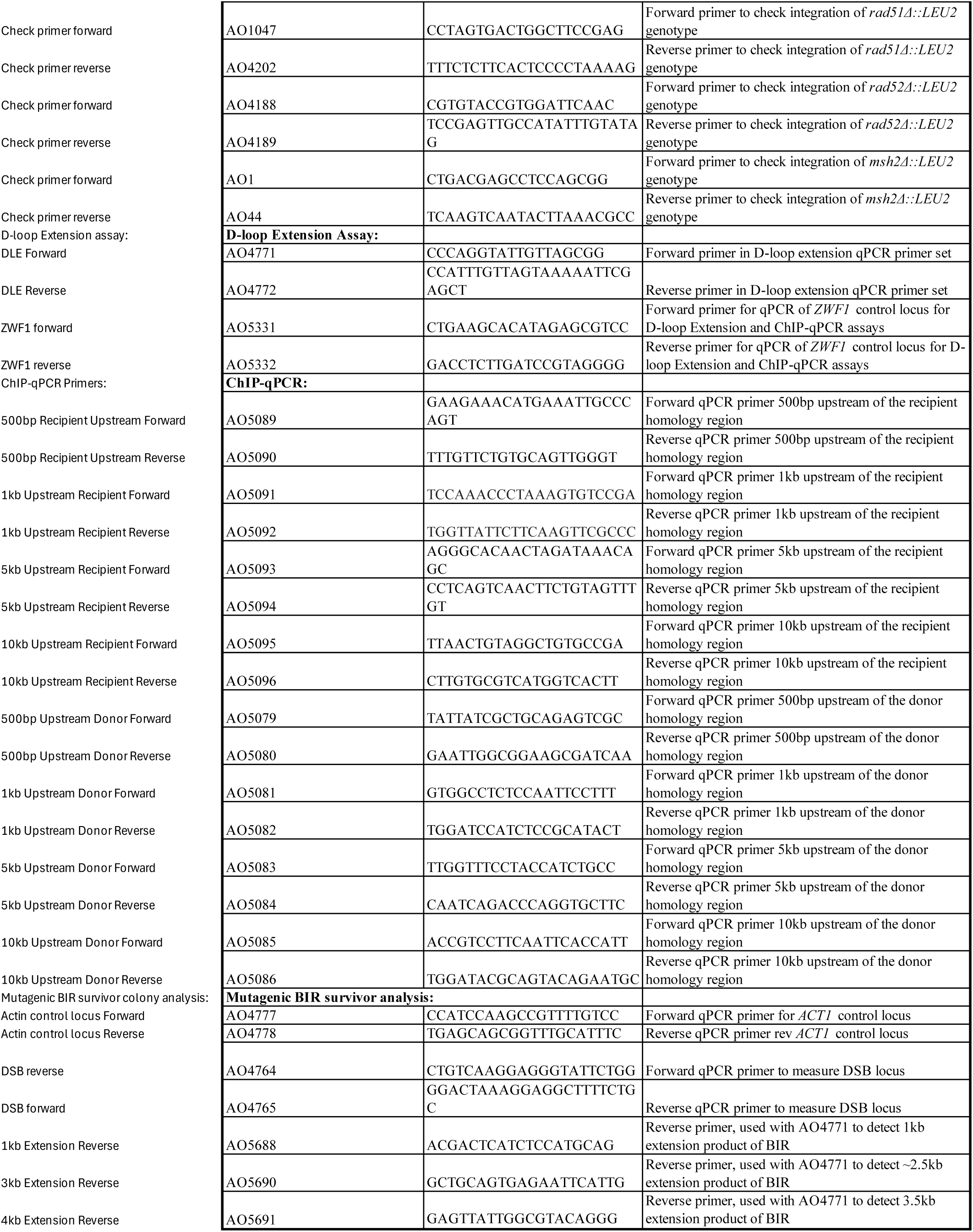
Oligonucleotides (shown 5’ to 3’) used in this study.

S1 Data. Underlying Data, Wild-type Ura^+^ survival assay.

S2 Data. Underlying Data, **Fig 1B**; 1C; 1D; **Fig 6**.

S3 Data. Underlying Data, **Fig 1B**; 1E; 1F; S2 Fig.

S4 Data. Underlying Data, **Fig 1E**; 1F.

S5 Data. Underlying Data, **Fig 2B**; 2C; 2E; 2F; S3A Fig; S3B Fig.

S6 Data. Underlying Data, **Fig 2D**; 2G; S3C Fig; S4 Fig.

S7 Data. Underlying Data, **Fig 7C**.

S8 Data. Underlying Data, **Fig 7E**.

S9 Data. Underlying Data, S1 Fig.

